# Evaluation of subretinally delivered Cas9 ribonucleoproteins in murine and porcine animal models highlights key considerations for therapeutic translation of genetic medicines

**DOI:** 10.1101/2024.12.30.630799

**Authors:** Spencer C. Wei, Aaron J. Cantor, Jack Walleshauser, Rina Mepani, Kory Melton, Ashil Bans, Prachi Khekare, Suhani Gupta, Jonathan Wang, Craig Soares, Radwan Kiwan, Jieun Lee, Shannon McCawley, Vihasi Jani, Weng In Leong, Pawan K. Shahi, Jean Chan, Pierre Boivin, Peter Otoupal, Bikash R. Pattnaik, David M. Gamm, Krishanu Saha, Benjamin G. Gowen, Mary Haak-Frendscho, Mary J. Janatpour, Adam P. Silverman

## Abstract

Genetic medicines, including CRISPR/Cas technologies, extend tremendous promise for addressing unmet medical need in inherited retinal disorders and other indications; however, there remain challenges for the development of therapeutics. Herein, we evaluate genome editing by engineered Cas9 ribonucleoproteins (eRNP) in vivo via subretinal administration using mouse and pig animal models. Subretinal administration of adenine base editor and double strand break-inducing Cas9 nuclease eRNPs mediate genome editing in both species. Editing occurs in retinal pigmented epithelium (RPE) and photoreceptor cells, with favorable tolerability in both species. Using transgenic reporter strains, we determine that editing primarily occurs close to the site of administration, within the bleb region associated with subretinal injection. Our results show that subretinal administration of eRNPs in mice mediates base editing of up to 12% of the total neural retina, with an average rate of 7% observed at the highest dose tested. In contrast, a substantially lower editing efficiency was observed in minipigs; even with direct quantification of only the treated region, a maximum base editing rate of 1.5%, with an average rate of <1%, was observed. Our data highlight the importance of species consideration in translational studies for genetic medicines targeting the eye and provide an example of a lack of translation between small and larger animal models in the context of subretinal administration of Cas9 eRNPs.

## Introduction

Cas9 endonucleases have emerged over the last decade as highly promising tools for in vivo editing of somatic cells to treat genetic diseases (Macarrón Palacios et al., 2024; Raguram et al., 2022; Taha et al., 2022). Cas9 gene editors use a guide RNA (gRNA) to direct the endonuclease to a DNA target site where Cas9 introduces a double stranded break (DSB). Correction of the DSB by intrinsic DNA repair mechanisms, predominantly non-homologous end joining (NHEJ), will frequently lead to insertions or deletions (indels) (Cong et al., 2013; Doudna and Charpentier, 2014). As indels typically cause nonsense mutations in the gene sequence, this approach enables knockout of a native gene. However, many genetic disorders are caused by point mutations that diminish the expression or function of a gene product and therefore cannot be corrected through introduction of indels. The need for an alternative approach to correct point mutations led to the development of base editors, which typically use a modified Cas9 nuclease that introduces a single-stranded nick rather than a DSB (nCas9), along with an engineered adenosine deaminase or cytosine deaminase that can respectively convert A to G or C to T (Anzalone et al., 2020; Gaudelli et al., 2017; Komor et al., 2016).

The ophthalmic disease space is an attractive area for applying these technologies (Choi et al., 2023; Sundaresan et al., 2023), as the eye is an immune-privileged site (Streilein, 2003) and safe, highly localized routes of direct administration to the retina have been adopted clinically (Davis et al., 2019). Inherited retinal disorders (IRDs) represent an area of high unmet need for patients and significant challenges for drug developers. IRDs can be caused by either loss-of-function mutations inherited in an autosomal recessive manner (e.g., Stargardt disease (Huang et al., 2022) and Usher syndrome (Toualbi et al., 2020)) or by toxic gain-of-function mutations that are inherited in an autosomal dominant manner (e.g., certain forms of Retinitis Pigmentosa (Daiger et al., 2013; Zhen et al., 2023)). Gene augmentation therapy for autosomal recessive IRDs by AAV-mediated delivery of a replacement copy of the dysfunctional gene is one potential avenue of treatment, as exemplified by the FDA approval of Voretigene neparvovec (Luxterna) for RPE65-mediated Leber congenital amaurosis in 2017 (Smalley, 2017). However, a specific challenge for gene replacement in genetic ocular diseases is that the large size of the cDNA encoding proteins involved in many IRDs far exceeds the packaging capacity of established delivery vectors (Ail et al., 2023); for example, the cDNA sizes for replacement of *ABCA4* (Stargardt disease) and *USH2A* (Usher syndrome) are 6,819 bp and 15,606 bp, respectively, and cannot fit within the ∼4.7 kbp packaging limit of AAV (Wu et al., 2010). Furthermore, gene augmentation cannot address autosomal dominant disorders.

Conversely, gene editing offers a potentially transformative treatment avenue for IRD patients, and the availability of different formats of gene editors could enable fit-for-purpose gene correction for different types of disorders: introduction of indels by NHEJ editors to address autosomal dominant IRDs and correction of point mutations by base editors to address autosomal recessive conditions. In order to bring gene editing therapies to patients, preclinical studies must demonstrate effective editing by the therapeutic in the cell types of interest in animal models with as much similarity to human physiology as possible. While mice have traditionally served as a preclinical model to assess efficacy and safety during drug development, mouse eyes have limited utility for evaluating drugs to treat IRDs due to anatomical differences with human eyes and the lack of models that accurately recapitulate human disease.

Therein lie two significant challenges in the development of gene editors as therapeutic agents: delivering functional gene editors to the nucleus of target cells, and translating findings from animal studies to humans during preclinical development. While AAV is the most frequently employed delivery vehicle for Cas9 gene editors, it suffers from several drawbacks including substantial immunogenicity (Bucher et al., 2021), potential to cause retinal atrophy (Reichel et al., 2023), and the risks of off-target edits due to long-term expression of the endonuclease (Doudna, 2020). Lipid nanoparticle (LNP) delivery of mRNA encoding for Cas9 gene editors is also being explored (Chien et al., 2022; Herrera-Barrera et al., 2023), but current LNP formulations show limited delivery to photoreceptors and safety concerns have been reported (Ana et al., 2022). Evaluating the delivery efficiency of therapeutic gene editors is further complicated by differences in the genetics of translational species. The therapeutic gRNA delivered with the Cas9 endonuclease specifically targets a human disease mutation; directly evaluating the therapeutic gRNA in a wild-type animal is usually impossible because (a) the animal does not have the disease mutation and (b) the DNA sequence of gene editing target loci may not be identical to human. The absence of a human transgenic disease model comprising the specific mutation of interest (few of which exist for IRDs), this necessitates using surrogate gRNAs, which in turn complicates evaluations of delivery efficiency because different gRNAs may inherently perform differently.

In this work, we describe a series of studies evaluating gene editors for ophthalmic diseases delivered as fully formed ribonucleoprotein (RNP) complexes administered with no delivery agents or encapsulation (Figure 1A). We characterize both i) NHEJ-eRNPs (double strand break inducing nuclease Cas9 constructs) and ii) BE-eRNPs (nickase Cas9-adenine base editor constructs). We explore the translational potential of this approach by delivering BE-eRNPs and NHEJ-eRNPs directly to the retinal pigmented epithelium and photoreceptors via subretinal injection in mice and Yucatan minipigs (Figure 1B), while screening sgRNAs in cell-based assays and mice to support the identification of translatable species-specific gRNAs (Figure 1C).

**Figure 1.**
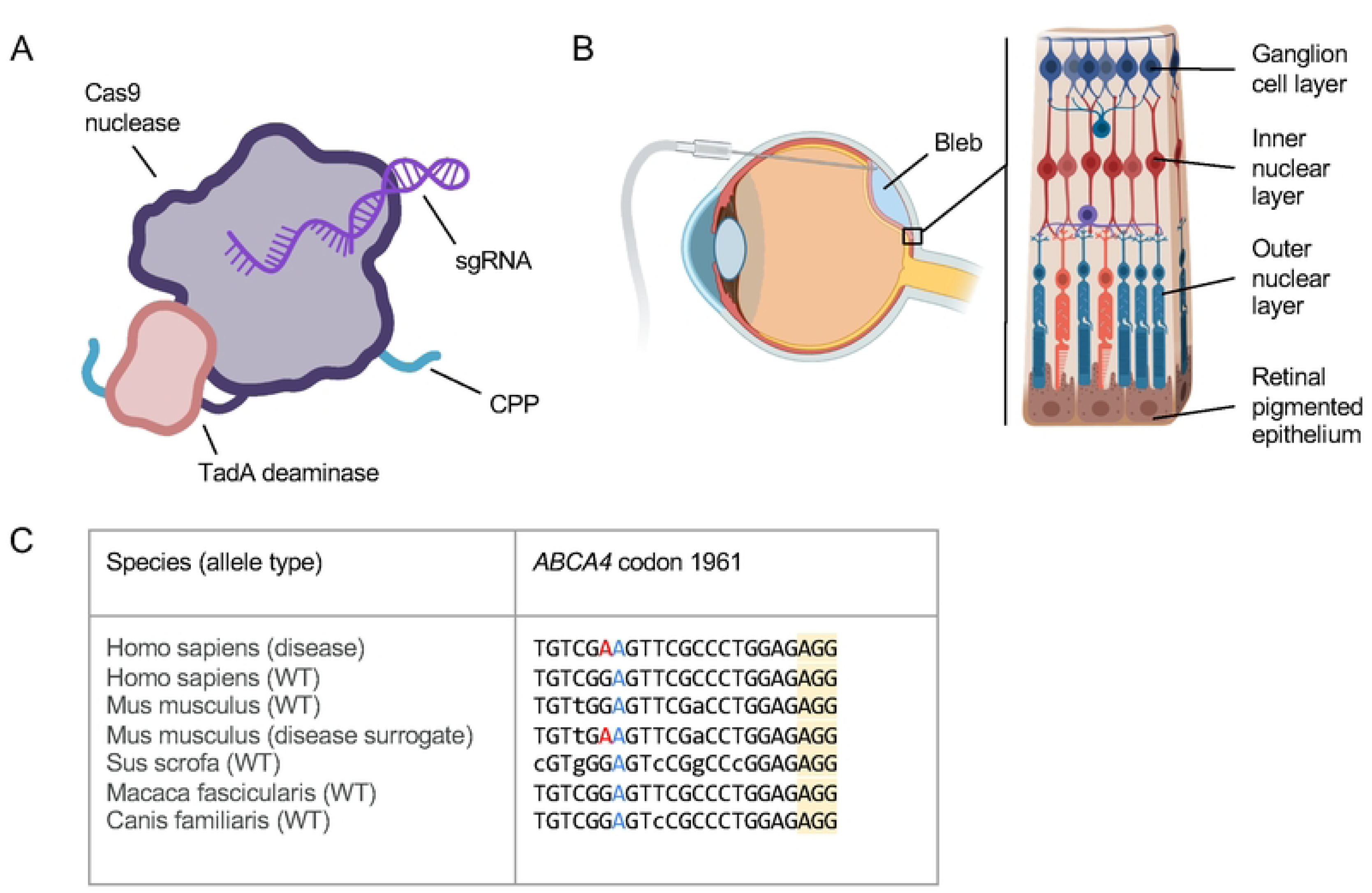
Overview of engineered ribonucleoprotein (eRNP) approach and challenges to treat inherited retinal disease. A. Features of adenine base editing engineered ribonucleoprotein (BE-eRNP). CPP, cell-penetrating peptide; sgRNA, single guide RNA. B. Subretinal delivery route and retinal anatomy. The bleb is formed between the outer nuclear layer containing photoreceptors and the retinal pigmented epithelium. C. Aligned genomic DNA sequences from the indicated species and alleles corresponding to one human disease variant underlying Stargardt’s Disease, *ABCA4* c.5882G>A, p.G1961E. The sequences shown correspond to a protospacer sequence and protospacer adjacent motif (PAM, yellow highlight) for WT SpCas9 with the disease-causing missense mutation (A in the human disease allele) colored red. A bystander A, colored blue, is present in all species shown but would not change the coding sequence of the gene if edited to G by an ABE. Lowercase bases indicate positions in the protospacer that vary relative to the human sequences.

## Results

### Generation and validation of NHEJ and ABE eRNPs

We based our nucleases on Cas9 from *Streptococcus pyogenes* Cas9 (SpCas9) and genetically fused SV40 nuclear localization sequences (NLS). The NHEJ-eRNP constructs include a SpCas9 lacking cysteines (C80S, C574S), while the BE-eRNP constructs comprise nCas9 (D10A, C80A) and a single copy of engineered *Escherichia coli* (*E. coli*) TadA variant from ABE8.20 (Gaudelli et al., 2020). We expressed the proteins in *E. coli* and purified them using several chromatography steps, and then combined them with sgRNA to form highly pure, low endotoxin, and homogenous eRNPs (see Materials & Methods). NHEJ eRNPs were strictly monomers and exhibit in vitro nuclease activity while BE-eRNPs formed dimers and exhibited in vitro deamination activity (Supp Figures 1–3).

**Figure 2.**
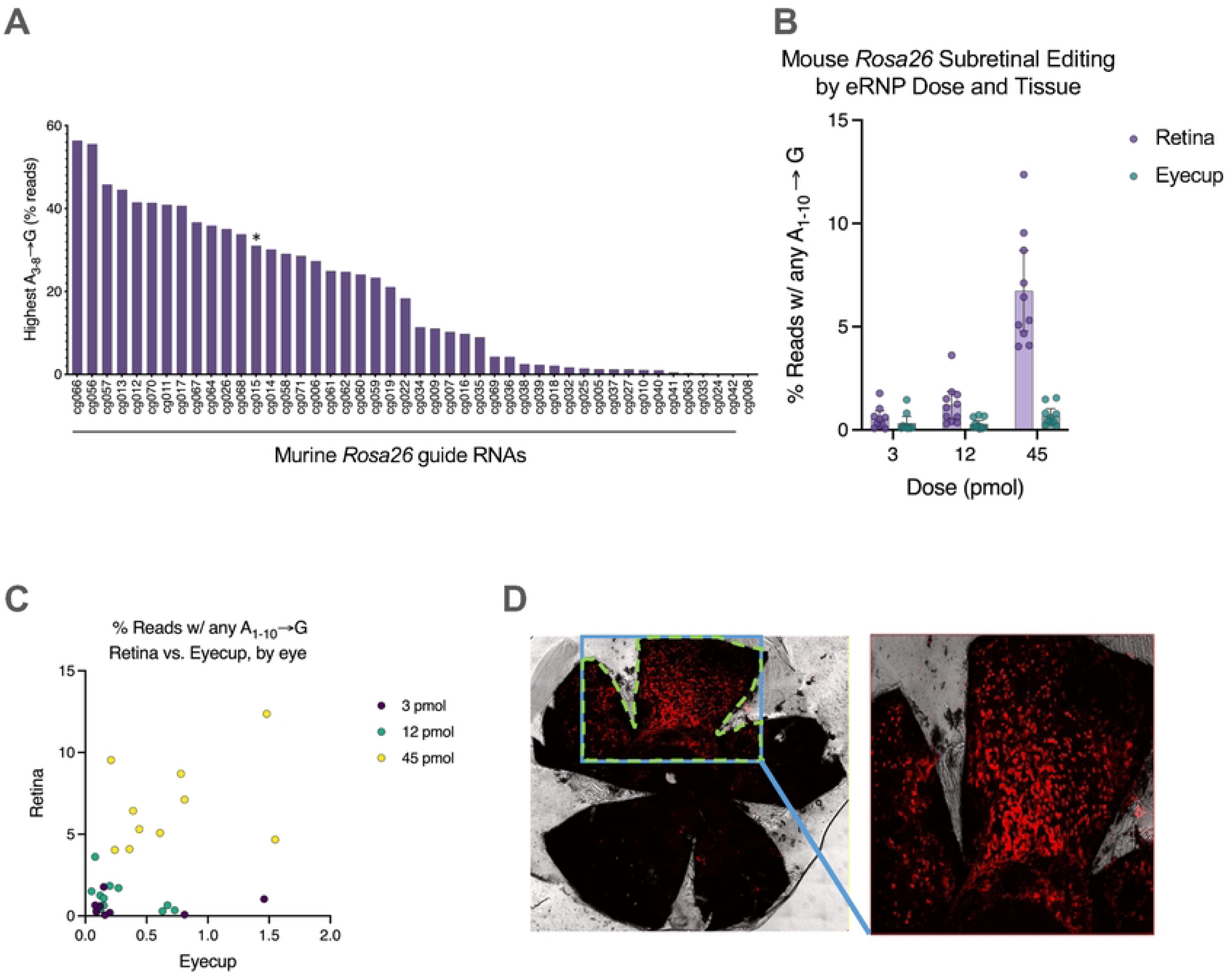
Identification of high efficiency ABE gRNAs targeting the mouse *Rosa26* locus and eRNP editing upon subretinal injection in mice. **A.** Editing rates in mouse T cells for different gRNAs delivered by ABE-eRNPs. ABE-eRNPs were complexed with the indicated gRNAs in crRNA:trcrRNA format and nucleofected into cells. Editing rates were quantified based on read frequency from Illumina sequencing and are displayed as averages per gRNA. The asterisk indicates the crRNA corresponding to the targeting gRNA used in panels (B) and (C). **B.** Editing rates by tissue following subretinal injection of Rosa26-targeting ABE-eRNPs complexed with mmRosa_sg1 in mice. 7 days following subretinal administration, mice were sacrificed and their eyes were dissected to separate the neural retina from the eyecup (containing choroid, sclera and RPE). Editing rates from Illumina DNA amplicon sequencing are shown as mean ± standard deviation (SD) at each dose. Each overlaid point corresponds to a single eye. Reads were scored as positive for editing if at least one A→G transition was detected within a 10-base editing window. **C.** Editing rates from the same samples in (B) plotted as editing in retina vs. eyecup tissues, colored by RNP dose. Each point corresponds to one eye. **D**. Fluorescent images of entire RPE tissue florets from Ai14 mice dissected from enucleated eyes 2 weeks after injection with NHEJ-eRNP complexed with sgAi14. The RPE florets were flat-mounted to assess genome editing by imaging the tdTomato (red) fluorescence through confocal microscopy.

**Figure 3.**
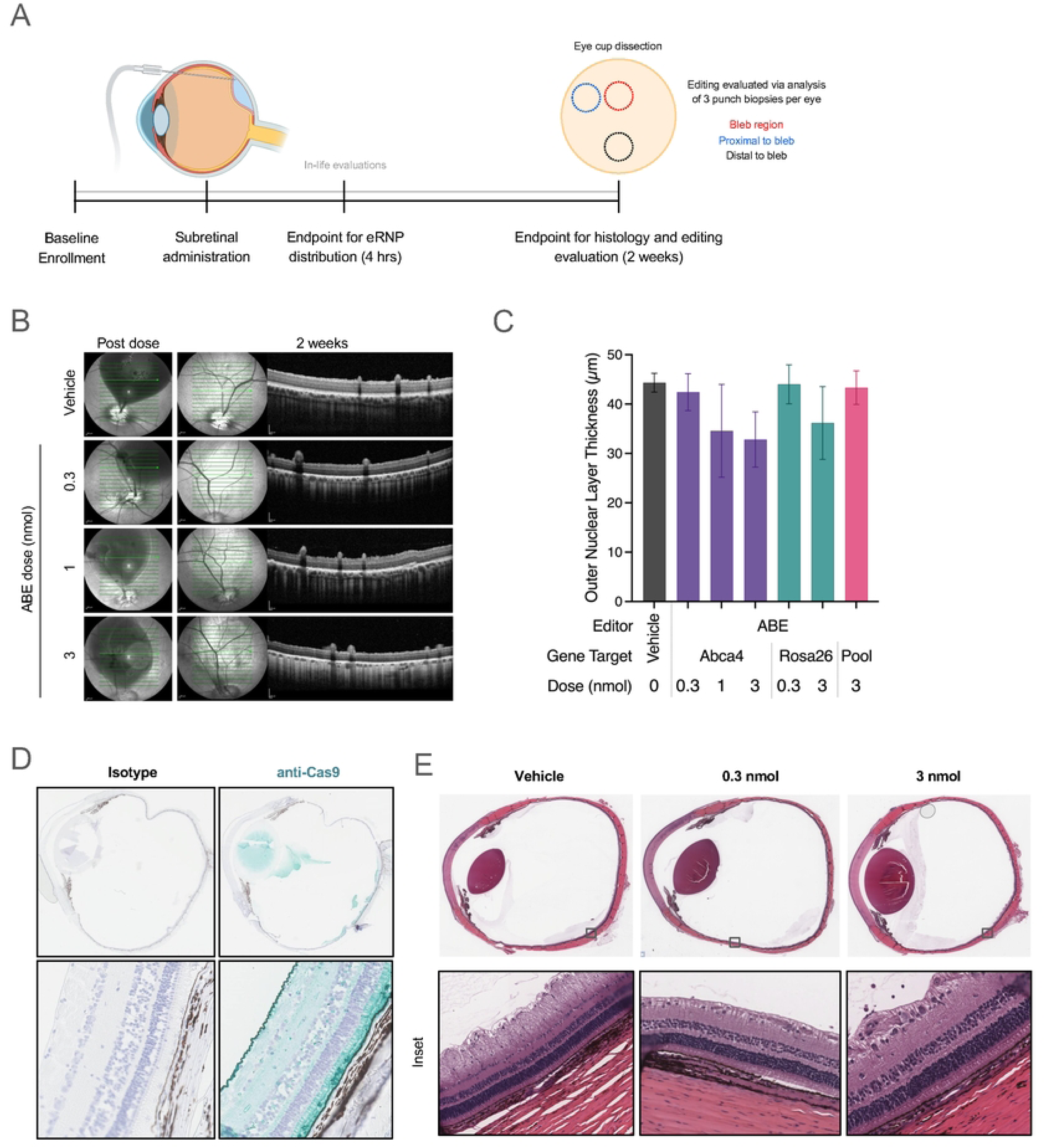
Physiology and histology of porcine eyes following subretinal administration. **A.** Graphical summary of the experimental design to evaluate the effects of subretinal administration of eRNP in Yucatan minipigs. **B.** OCT images of porcine eyes immediately following and 2 weeks following subretinal dosing of vehicle or ABE-eRNPs at the indicated dosages (b-scan indicated by green arrow). **C.** Outer nuclear layer thickness according to OCT at 2 weeks post injection plotted for ABE-eRNP groups. Data are plotted as the mean ± SD for each group across all measurement areas from superior (injected) regions, as shown for representative eyes in (A). **D.** Representative images of porcine eyes 4 hours post subretinal administration stained with isotype control or anti-Cas9 immunohistochemistry (teal) at 2.5× (top) and 20× (bottom) magnification. **E**. Representative whole eye scans and high magnification (inset) images of H&E-stained porcine eyes 2 weeks post subretinal administration with either vehicle, 0.3 nmol ABE-eRNPl or 3 nmol ABE-eRNP.

### Guide RNA identification and selection

To evaluate the eRNPs as a therapeutic modality for the treatment of IRDs, we aimed to measure their potency when administered via subretinal injection. To enable these studies, we first set out to identified high-efficiency gRNAs that enabled evaluation of base editing and NHEJ. We designed 71 sgRNAs targeting the murine *Rosa26* locus, informed by previously reported sequences (Abe et al., 2020; Chu et al., 2016; Lusk et al., 2022; Quadros et al., 2015) (Supp. Table 2). We evaluated the NHEJ and ABE activity of these sgRNAs by nucleofection of primary mouse T cells. The editing efficiency of these gRNAs varied substantially, with ABE editing efficiency ranging from 0% to greater than 55% (Fig 2A) and NHEJ editing efficiency ranging from 5% to 60% (Supp Figure 4). Interestingly, NHEJ and ABE editing rates of individual gRNAs correlated (Supp Fig 4B). Together, these data identify high efficiency gRNAs targeting the murine *Rosa26* locus.

**Figure 4.**
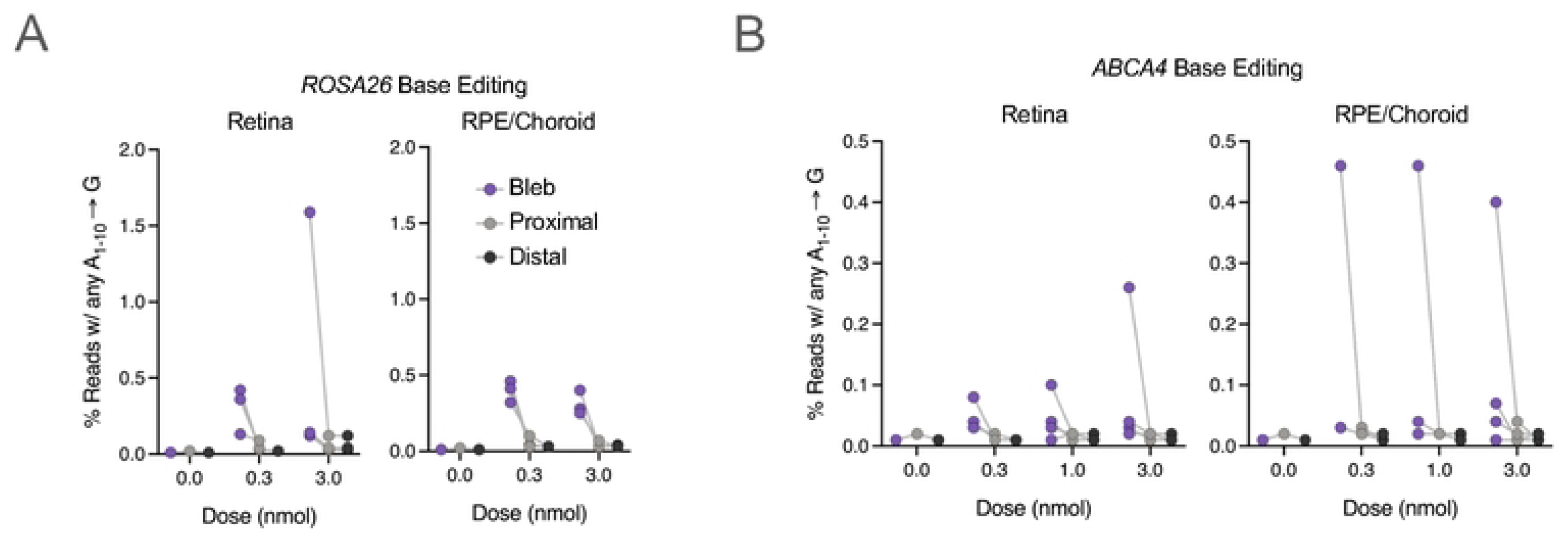
Adenine base editing following ABE-eRNP subretinal administration in minipig eyes. **A.** Editing rates by tissue following subretinal injection of *ROSA26*-targeting ABE-eRNPs complexed with sgRosa26 in minipigs at multiple doses. Two weeks following subretinal administration, animals were sacrificed, and eyes were dissected to collect intact tissue layers for the neural retina and the choroid + retinal pigmented epithelium (RPE). For each tissue, three biopsy punches were taken corresponding to the injection bleb, tissue immediately adjacent to the bleb, and a region distal to the bleb. Editing rates from Illumina DNA amplicon sequencing are plotted for each eye by tissue, biopsy region, and dose. Reads were scored as positive for editing if at least one A→G transition was detected within a 10 base edit window. Gray lines connect points corresponding to one eye. **B.** Editing rates as shown in (A) for ABE-eRNPs complexed with sgABCA4.

### Subretinal administration of eRNPs in mice

To assess what cell types are exposed to eRNPs following subretinal administration, we provided a single subretinal injection of an eRNP to mice and sacrificed the animals at 10 minutes, 4 hours, or 24 hours after dosing. Following enucleation, we sectioned the eyes to determine intraocular distribution of the eRNP using anti-Cas9 immunohistochemistry (IHC). All eyes showed retinal detachment, which was most pronounced at the 10-minute time point, consistent with the formation of a subretinal bleb. After staining sections for Cas9 protein, we observed IHC signal within the subretinal bleb region, primarily in the outer nuclear layer of photoreceptor cells (Supp Fig. 5A). The pigmentation of the retinal pigmented epithelial (RPE) cells prevented clear interpretation of the Cas9 signal, though detection of editing by next-generation sequencing (NGS, see below) indicates that these cells also took up RNP.

**Figure 5.**
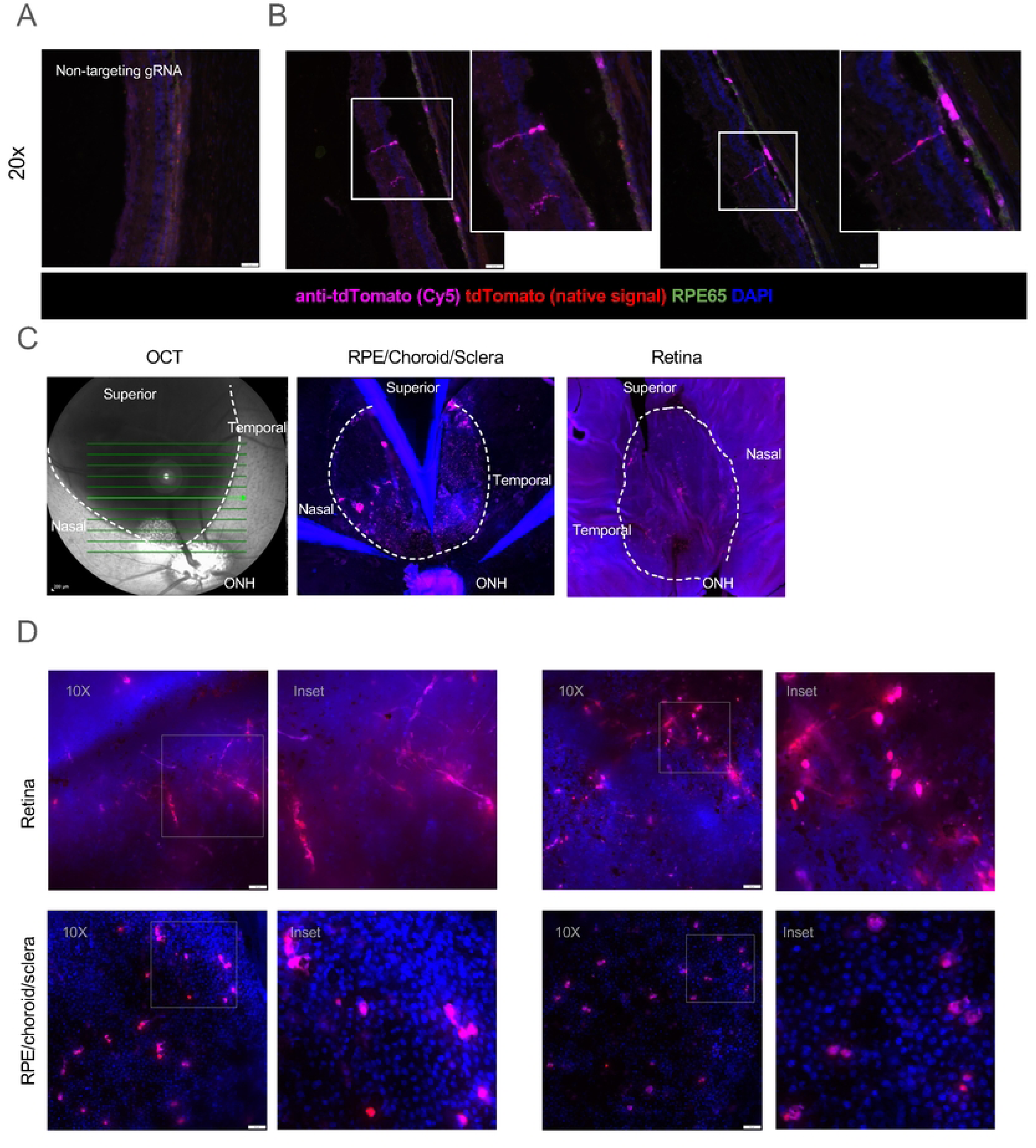
Characterization of editing in the transgenic SRM1 pig strain. Pseudocolored images show staining by DAPI (blue), native tdTomato (red), anti-tdTomato-Cy5 (magenta) **A.** Fluorescent image of cryosection of an SRM1 transgenic porcine eye treated with a negative control non-targeting gRNA NHEJ-eRNP. RPE65 is co-stained for RPE visualization (green). **B**. Fluorescent images of the superior (treated) region of the two SRM1 transgenic porcine eyes subretinally administered targeting gRNA NHEJ-eRNP. Scale bar, 50 microns. Insets show higher magnification of the boxed regions. **C**. OCT (left) and fluorescent images of flat-mounted tissue (middle and right) after subretinal administration of eRNP. The bleb region is outlined with a dotted line with anatomical regions labeled. **D**. Fluorescent images of retina and RPE/choroid/sclera flat-mounted tissue for eyes from the two SRM1 pigs administered targeting gRNA NHEJ-eRNP. Insets show higher magnification of the boxed regions.

We then assessed the editing activity of eRNPs upon subretinal administration in C57BL/6 wild-type mice. BE-eRNPs were administered via subretinal injection and we confirmed bleb formation by post-dose optical coherence tomography (OCT) imaging. Ocular examinations did not reveal any significant safety findings associated with test article administration (Supp Table 1). We assessed editing rates in the retina and remaining eye cup 7 days after dosing by isolating and sequencing genomic DNA. We observed a dose-response for BE-eRNPs, with highest editing at the 45 µM dose and an average of ∼7.5% editing in retina tissue (Fig 2B). Interestingly, base editing rates were significantly higher in retina compared to RPE with a moderate correlation between editing rates observed in retina and eyecup tissue samples from individual eyes (Fig 2C).

To assess the biological and technical reproducibility of these results we repeated the mouse subretinal injection at a different institution, utilizing the identical lot of BE-eRNP material as the first study (Supp. Fig. 5C). The overall findings were consistent with the first study, with a trend towards lower absolute editing levels in the retina, a similar dose response, and an observation that ABE editing with BE-eRNP is higher in the retina than the RPE. In this study, we isolated the RPE from the eye cup and performed NGS analysis on gDNA from the RPE alone, rather than the entire RPE-choroid layer. As such, the editing rates were higher in RPE in this study as compared to the RPE/choroid editing levels in the initial study, consistent with the notion that RPE are the predominant cell type edited within the RPE/choroid eye cup sample.

We next evaluated the distribution of editing via imaging of RPE flat-mounts upon subretinal administration in Ai14 reporter mice, in which editing induces TdTomato expression. At day 14 following administration, eRNP-treated mice showed bright tdTomato signal in the RPE floret, covering 5–25% (average 12.02 ± 1.6%) of the floret area (Fig 2D and Supp Fig 5D).

Collectively, these data suggest that eRNPs can efficiently edit their target cells in mice when delivered via subretinal administration. Two factors that may impact observed editing rates (1) the observation that Cas9 is only exposed to a fraction of the retina, likely less than half with a 1 µl injection volume (Supp Fig 5B), and (2) that the editing was assessed based on the all cells within the retina or RPE (or RPE/choroid), Based on these observations, we hypothesized that localized editing within the bleb region was significantly higher than the editing rates measured for the entire eye. The size of the mouse eye precludes microdissection to evaluate localized editing within the bleb, but such an analysis is possible in larger animals.

### Subretinal administration of RNPs in minipigs

Given the similarities in size and physiology between human and porcine eyes, pigs are considered a useful translational model for evaluating ophthalmic drugs that minimizes the use of non-human primates. We next evaluated the activity of subretinally administered eRNPs in minipigs, in which we could also evaluate gene editing specifically at the site of administration as well as regions adjacent or distal to the bleb. We first identified sgRNAs targeting the *S. scrofa Rosa26* locus, which is poorly conserved between *M. musculus* and *S. scrofa.* We selected two sgRNAs for evaluation: a previously reported sgRNA with validated NHEJ activity (Li et al., 2017) and an sgRNA targeting porcine *ABCA4* at the site homologous to human disease allele *ABCA4* c.5882G>A, p.(G1961E) (Supp Fig. 6A, Supp Table 2). The *Rosa26*- and *ABCA4*-targeting gRNAs were active in two porcine cell lines (Supp Fig. 6A).

In consideration of the fact that in vivo editing activity is dependent on sgRNA, and that cell-based characterization may not be predictive of in vivo activity, we performed an in vivo pooled sgRNA screen to evaluate the performance of a larger number of sgRNA sequences than could reasonably be tested individually. We complexed ABE protein individually with nine different sgRNAs, and then combined them at an equimolar ratio to generate pooled RNP test article. Following subretinal administration of 3 nmol of the pooled BE-eRNP there were no significant adverse findings observed by ocular examinations. We observed editing with each of the sgRNAs, however, the dynamic range was not sufficient to rely on this approach in its current form for evaluation of therapeutic potential of eRNPs (Supp Fig. 6B). As such we selected one gRNA targeting ssRosa26 and one targeting ssAbca4 to evaluate NHEJ- and BE-eRNPs in Yucatan minipigs single gRNAs, based on the in vitro cell based assay data.

The goals of our study were multifold: (a) determine whether NHEJ and/or BE-RNPs can potentially achieve clinically meaningful editing rates upon subretinal administration, (b) assess eRNP distribution and editing rate inside and outside of the bleb, (c) pursue dose finding to inform higher powered future studies, and (d) determine any test article-mediated safety signals with research-grade material. We generated ABE and NHEJ eRNPs to meet stringent specifications of purity, in vitro activity, and endotoxin levels and confirmed the compatibility of the test articles with two injection devices (MedOne and DORC). Our study design is shown in Figure 3A. The subretinal dose was targeted to the area centralis at midline in the superior retina. For subsequent imaging and histological examination, the inferior retina was presumed to be uninvolved. OCT and color fundus imaging performed immediately after dosing confirmed bleb formation.

In-life ocular examinations were generally unremarkable, with minor acute inflammation observed at day 3 that resolved by day 7 in all groups, including vehicle. OCT and fundus photography 2 weeks post-dosing showed no major findings across all dose levels of BE-eRNPs, including no evidence of significant inflammation or gross retinal degeneration (Fig 3B, Supp Fig 6C). Evidence of potential outer segment (OS) loss, but no nuclear layer loss, was consistent with incomplete healing and OS restoration at 2 weeks post-dose. In contrast to the observed tolerability of BE-eRNPs at the highest dose of 3 nmol, we observed outer nuclear layer (ONL) thinning in the superior retina with the 3 nmol dose of NHEJ eRNP (Fig 3C, Supp Fig 6D-E). This appeared to be a dose-dependent effect, as no ONL thinning was observed at a 0.3 nmol dose of the same test article.

We determined the intraocular distribution of the test articles by anti-Cas9 IHC on eyes that were fixed 4 hours post-dosing. The Cas9 signal was largely restricted to RPE and outer segments within an area of the superior retina, consistent with the subretinal bleb, while Cas9 was not detected in the inferior retina (Fig 3D). Cas9 was not detected in the RPE or outer segments 2 weeks following eRNP administration, with limited IHC signal observed on the surface of the retina (Supp Fig 6F). Histopathology indicated that the eRNP test articles were well tolerated with minimal findings observed in most evaluated eyes (Figure 3E). We observed slight mononuclear cell infiltrate in the vitreous in all groups, including the vehicle group, indicating mild inflammation. One eye had qualitatively different inflammatory cell infiltrate at the pars plana sclerotomy site, which was assessed to be procedure related. Consistent with the OCT findings, we saw diffuse superior outer retinal degeneration with the high dose of NHEJ eRNP (Supp Fig 6G–H).

For each eye not used in histopathological assessments, we collected three 8 mm punch biopsies: within the bleb, proximal to the bleb, and distal to the bleb (See Methods). The punch biopsy was estimated to capture approximately 90% of the bleb region based on biopsy diameter and approximate bleb size. We separated the neural retina from the RPE/choroid in each punch, and then extracted genomic DNA to assess editing levels by NGS. Although we observed editing in both NHEJ- and BE-RNP groups in both RPE/Choroid and retinal tissue samples, no samples showed greater than 2% indel formation or A→G transversion, and most samples showed <0.5% editing efficiency (Fig. 4A, 4B, and Supp Fig 7A, 7B). All editing outcomes observed were consistent with test article and gRNA identity. There did not appear to be a dose-response in the BE-eRNP samples tested at 0.3, 1, and 3 nmol. There was no editing observed in vehicle controls or in uninvolved loci (i.e., *ABCA4* editing for *Rosa26*-targeted RNPs or vice versa). The greatest editing was in the bleb regions, consistent with limited distribution of eRNPs beyond the injection site, though some editing was also seen in the regions proximal to the bleb (Fig 4A, 4B).

Because *ABCA4* is only expressed in a subset of cells within the eye (primarily in photoreceptors, but to a lesser extent in RPE), we assessed the editing rate at the mRNA level via cDNA sequencing for two eyes that were treated with an *ssABCA4* BE-eRNP (receiving high dose and medium dose). No significant difference in editing rates via sequencing of gDNA vs cDNA were observed (Fig 4B, Supp Fig 7A). In one of two eyes, a higher editing rate in the RPE was observed via cDNA sequencing (0.3 %) compared to gDNA sequencing (near limit of detection). This raises the possibility that editing events may occur at a higher rate in ABCA4 expressing cells (RPE) than non-ABCA4 expressing cells (e.g., cells in the choroid) within the RPE/choroid sample; however, this hypothesis requires additional investigation.

### Subretinal administration of RNPs in transgenic SRM1 reporter pigs

To determine which cell types and anatomical sites are edited following subretinal administration of eRNPs, we utilized a porcine transgenic reporter strain in which Cas9 mediated NHEJ editing leads to expression of tdTomato (See methods). We confirmed the catalytic activity of the eRNP for this target locus (Supp Fig 8A). Based on our prior observation of ONL thinning associated with subretinal administration of 30 µM NHEJ-eRNP, but not BE-eRNP, we selected a dose of 15 µM. Minor reduction in ONL thickness was observed (Supp Fig. 8B) and ocular examinations did not reveal major findings (Supp Fig. 8C).

We assessed the fidelity of fluorescent signal in this pig reporter model via immunofluorescent imaging of cryo-sections. Minimal tdTomato signal was observed in littermate wild-type control pigs administered with vehicle or targeting eRNPs, as well as in transgenic animals administered with eRNPs comprising non-targeting gRNA (Fig 5A, Supp Fig. 8C). Extensive controls for background fluorescent signal in various regions are shown in Supp Figure 8; in general, some background was observed in the native tdTomato channel, with less in samples stained with labeled secondary antibodies. In the transgenic reporter pig administered eRNP with targeting gRNA, we observed tdTomato fluorescent signal within the treated (superior) region via both direct and indirect fluorescent detection in one of the two eyes assessed (Fig 5B and 5C). The anatomical location and morphology of fluorescent signal suggested that RPE cells and Müller glia were the primary cells edited. These findings support the fidelity of the SRM1 transgenic reporter model and indicate that subretinal administration of eRNP can mediate editing of specific cell types within the neural retina and RPE.

We next assessed tdTomato fluorescence of flat mounted retina and RPE/choroid/sclera tissue to determine the distribution of edited cells. Comparison of OCT images captured immediately after dosing with flat-mount fluorescent imaging data indicates that editing primarily occurred within the region of the subretinal bleb (Fig 5C). We observed tdTomato signal in both retina and RPE flat-mounts of transgenic tissue (Fig 5D). Fluorescent signal was not observed in wild-type tissue nor in the untreated inferior region of the transgenic eye, further supportive of the fidelity of the model (Supp Fig 8F). tdTomato fluorescence was observed in neuronal projections in the retina and in RPE cells (Figure 5E, insets).

## Discussion

Herein, we report studies in mice and pigs conducted to support therapeutic development of an in vivo Cas9 genome editing platform. In the context of retinal disorders, while small animals can provide useful initial insights into the activity of therapeutic candidates, large animal models more closely recapitulate the anatomy and physiology of the human eye (e.g., globe size, foveation/photoreceptor distribution and density) (Chader, 2002; Winkler et al., 2020). Direct in vivo administration of eRNPs is fundamentally distinct from AAV or LNP based approaches. We demonstrated that subretinal administration of Cas9 eRNPs can mediate in vivo editing in mice and pigs, albeit with differential activities. We observed differences in the editing outcomes in the pig and mouse animal studies, which complicate therapeutic development and may inform the selection of preclinical animal models for the evaluation of similar genetic medicines in the eye. Direct comparison of editing rates in mice and pigs is difficult because of intrinsic differences in the assessment methodologies enabled for each model. In mice, it is not technically feasible to isolate the treated region, and as such the observed editing rates reflect editing efficiency with the entire retina or eyecup. In contrast, the size of the pig eye enables dissection of the treated bleb region only. As such, with an assumption that editing primarily occurs within the treated bleb region (supported by our reporter animal studies) and that editing was observed primarily within the treated bleb region in our pig studies, it is reasonable hypothesize that the editing rates observed in the mouse studies were underestimates of the editing rate within the treated region (perhaps by 2–3-fold based on the Cas9 distribution and RPE flat-mount imaging of Ai14 reporter mice).

The observed differences between genome editing rates mediated by BE-eRNPs and NHEJ-eRNPs observed in mouse and pig models of subretinal administration highlights potential challenges for the development of genomic medicines to treat ophthalmic and other diseases. Potential considerations, depending on the specific therapeutic modality and disease indication include: i) conservation of the genomic target sequence (e.g., if use of a surrogate gRNA is necessitated), ii) use of rodent animal models (and/or cell models) as predictive models for translation, iii) technical aspects of route of administration (including injection method and apparatus differences), iv) anatomical differences in the target organ between small and large animal species, v) biological differences in cell types of interest between small and larger animal species and vi) differential mechanisms of eRNP absorption and clearance from the subretinal space (e.g., rate of elimination of eRNP and/or subretinal bleb volume via RPE).

Although porcine and human eyes have substantial anatomical differences (e.g., area centralis in the former rather than a macula in the latter), porcine eyes are similar to human eyes in axial length and size, providing a tractable large animal model. Several advantages to assess the translational feasibility include (1) that subretinal injection procedure can be performed in a clinically relevant fashion (similar volumes, consumables, apparatus), (2) the eye size is similar (though vitreous volume is different), and (3) functional and molecular assessment of activity within the subretinal bleb is more technically feasible. A working hypothesis, therefore, is that the activity in larger animals, such as pig, may more closely recapitulate activity in human. For example, large animal models have been previously utilized for the evaluation and development of AAV-based gene therapy for Leber Congenital Amaurosis (LCA) (Acland et al., 2005, 2001; Bennicelli et al., 2008). On the other hand, a significant challenge for therapeutic development is the limited number of test articles that can reasonably be assessed in larger animal models and their relatively higher cost. Our findings highlight key considerations when designing preclinical studies aimed at supporting therapeutic development of ophthalmic genetic medicines.

## Materials and Methods

### Protein Constructs

The ABE used for this study was based on ABEmax (Koblan et al., 2018) and the 8.20-m variant of ABE8 (Gaudelli et al., 2020). The expression construct consisted, in order from N- to C-terminus, of a His-tag (removed by HRV 3C protease during purification), a bipartite NLS sequence (Koblan et al., 2018), the TadA adenosine deaminase corresponding to the 8.20-m variant of ABE8 (Gaudelli et al., 2020), a 32-amino acid glycine/serine-rich linker, S. pyogenes Cas9 with D10A and C80A mutations, and another bipartite NLS sequence at the C-terminus.

The expression construct used for all NHEJ editing in this study except for the Ai14 reporter mouse experiment consisted of an N-terminal His tag, MBP tag (removed by HRV 3C protease during purification), S. pyogenes Cas9 with C80S and C574S mutations, 2 copies of the SV40 NLS sequence, a SpyCatcher002 bioconjugation tag (Keeble et al., 2017) (not used in this study), and finally 4 copies of the SV40 NLS sequence at the C-terminus.

For the Ai14 reporter mouse experiment of Fig 2D and Supp Fig 5D, the expression construct consisted of an N-terminal His tag (non-cleavable) followed by the HIV Tat^48–57^ cell penetrating peptide sequence, one copy of the SV40 NLS sequence, SpCas9 with a C80A mutation, and 2 copies of the SV40 NLS at the C-terminus.

Cas9 constructs were codon optimized for E. coli expression using a custom workflow based on the DNA Chisel python package (https://edinburgh-genome-foundry.github.io/DnaChisel) and expressed from a T5-lac promoter in a plasmid containing a kanamycin resistance marker and a pUC origin of replication.

### Protein Production

Proteins were expressed in E. coli strain BL21(DE3) in shake flasks using Terrific Broth Complete medium (Teknova) and overnight induction with IPTG at 18 °C. Proteins were purified by Ni^2+^ affinity, cation exchange, and size exclusion chromatography. Endotoxin was removed during the Ni^2+^ affinity by extensively washing the column with buffer containing 0.2 % w/v Triton X-114. The N-terminal affinity tags were removed by HRV 3C protease after the Ni^2+^ affinity step. After purification, the proteins were concentrated to 10-20 mg/ml in one of two buffers and stored at - 80 °C before their final formulation for studies. Protein prepared for mouse studies and cell-based assays was stored in 20 mM L-histidine, 100 mM L-arginine, 200 mM sodium chloride, and 5 % w/v sucrose, pH 7.5. Protein prepared for pig studies was stored in 50 mM HEPES, 100 mM sodium chloride, 10 mM sodium phosphate, and 1 % w/v sucrose, pH 7.3.

### RNP Formulation

For cell-based assays and the mouse subretinal injection experiments in Ai14 mice, RNPs were formulated in 20 mM L-histidine, 100 mM L-arginine, 200 mM sodium chloride, and 5 % w/v sucrose, pH 7.5. For all other mouse and pig studies, the RNPs formulated in a buffer containing 50 mM HEPES, 100 mM sodium chloride, 10 mM sodium phosphate, and 1 % w/v sucrose, pH 7.3.

For all but the gRNA screening experiments, a 100 nt sgRNA modified at nucleotides 1-3 and 97-99 with 2’O-methyl and phosphorothioate was chemically synthesized by Integrated DNA Technologies, Inc. (Coralville, Iowa). gRNA sequences are described in Supplemental Table 2. Lyophilized gRNA was resuspended either in a low-salt storage buffer containing 5 mM HEPES, 0.1 mM EDTA, pH 7.5 (cell-based assays and mouse studies) or RNP formulation buffer (pig studies) and stored at −80 °C. Just prior to mixing with nuclease, the gRNA was thawed on ice, diluted in formulation buffer and heated to 70 °C for 5 minutes. The gRNA was allowed to slowly cool at room temperature on the bench top to refold, then stored on ice. The same refolding procedure was used for the gRNA screening experiments using crRNA and tracrRNA, but RNA components (Integrated DNA Technologies, Inc.) were mixed at a 1:1 molar ratio prior to RNA refolding. gRNA and Cas9 were mixed together on ice at a 1.2:1 molar ratio of gRNA:Cas. Cas9 was added to formulation buffer, followed by slow addition of and gentle pipetting up and down to thoroughly mix. The reactions were incubated for 10 minutes at 37 °C, then cooled ice. RNP samples for animal studies were applied to a 0.2 µm hydrophilic PVDF centrifugal filter (Millipore) to remove insoluble fraction prior to storage at −80 °C. For animal studies, RNPs were assembled at a 10 % higher concentration than required for dosing to account for losses during assembly and filtration. The protein concentration within the RNP was measured after assembly and filtration using a Bradford or BCA assay with a standard curve generated from the purified protein at the concentration established by A280 and the calculated extinction coefficient. After filtration, RNP was diluted with formulation buffer to the final injection concentration if necessary. Final RNP concentrations indicated in the figures were based on Cas9 protein concentration, assuming that in the presence of excess gRNA, 100% of the Cas9 protein was incorporated into RNP. For in vivo test articles, samples were analyzed for endotoxin with Endosafe PTS Cartridges (Charles River Laboratories, Wilmington, MA).

### In vitro DNA cleavage assay

A FRET assay based on a previously reported method was adopted (Seamon et al., 2018). dsDNA molecules were synthesized with 5’-Cy3 and 3’-Cy5 ends. Reactions were initiated by mixing eRNPs with labeled dsDNA at 37 °C for 30 min in a reaction buffer containing 10 mM L-histidine, 100 mM L-arginine, 200 mM NaCl, 5% (w/v) sucrose, 1 mM MgCl_2_, pH 7.5. The reactions were quenched with an equal volume of 8 M guanidine-HCl and analyzed with a ClarioStar fluorescence microplate reader (BMG Biotechnology).

### In vitro DNA adenosine deamination assay

eRNP and a linearized plasmid substrate were incubated at a 10:1 molar ratio at 37 °C for 45 min in a reaction buffer consisting of 10 mM L-histidine, 100 mM L-arginine, 200 mM NaCl, 5 % w/v Sucrose, 1 mM MgCl_2_, 0.1 mM ZnCl_2_, pH 7.5. The reaction was quenched by placing on ice and adding 0.5 mM EDTA and 2.4 mU/mL proteinase K, followed by incubating at 37 °C for 15 min and then at 65 °C for 10 min. The plasmid DNA substrate was treated with 0.2 U/µL endonuclease V (New England Biolabs), which cleaves sites with inosine bases created by the TadA-Cas9. The reaction was incubated at 1 hour at 37 °C followed by 5 min at 65 °C, then quenched by adding 0.75 mg/mL proteinase K and incubating for 15 min at 50 °C. DNA fragments were separated with a Fragment Analyzer (Agilent Technologies), and the cut and uncut DNA bands were integrated with the ProSize software.

### Nucleic acid extraction and sequencing of porcine PBMCs

Blood was collected in EDTA coated tubes (BD) and extracted using DNeasy Blood & Tissue kits (Qiagen). Once eluted in nuclease free water, purity (A260/280) and concentration (ng/µl) were measured using a NanoDrop One (Fisher). After extraction, genomic regions around each gRNA were PCR amplified for sanger sequencing. Primers were designed with primer3 to amplify a 300-500 bp region around each gRNA. A PCR reaction was performed with 50 µl of Q5 Hot-Start High-Fidelity 2X Master Mix (New England Biolabs, M0494) both primers at a final concentration of 0.5 µM, 400 nanograms of extracted genomic DNA, and nuclease free water to a final volume of 100 microliters. PCR reactions were purified with a Monarch PCR & DNA Cleanup Kit (New England Biolabs, T1030) and spot checked for purity via gel electrophoresis. Samples were sent to Azenta for sanger sequencing with 1 ng/µl of purified product and 25 pmol primer in 15 µl reactions. The resulting ab1 files were analyzed for sequence homology to their corresponding genomic regions in the sus scrofa genome (susScr11) using Benchling software.

### Guide identification and screening

The ssABCA4 gRNA was designed for adenine base editing of the porcine *ABCA4* gene at position G1960. This gRNA is the porcine surrogate of a previously reported gRNA targeting human ABCA4 c.5882G>A, p.(G1961E) (Muller et al., 2023). Li and colleagues previously described ssROSA26 gRNA for NHEJ two of which showed higher activity and contain an adenine in the optimal ABE window for our construct (ROSA26-sgRNA R5 and R6 of Li et al., 2017). To identify additional gRNAs, the porcine and mouse ROSA26 loci were aligned. From the aligning regions, gRNAs were identified with an on-target score >50 (Doench et al., 2016) and the presence of adenine in positions A6 and/or A7. gRNA target sites were at least 400 bp apart to avoid interference in editing, amplification, or measurement from the presence of multiple *ROSA26* gRNAs in the pooled eRNP sample. gRNAs were validated for ABE activity by nucleofection of eRNP-BE into PK(15) and PT-K75 cells.

Activity of murine gRNAs in primary mouse cells were evaluated in line with previously reported methods (Gowen et al., 2024). Briefly, murine primary T cells were magnetically isolated from spleens from Balb/c mice (Charles River Laboratories) and were activated with 60 U/mL of recombinant mouse IL-2 (R&D Systems) and Dynabeads Mouse T-activator CD3/CD28 (Gibco). T cells were cultured at 37°C in a humidified atmosphere with 5% CO_2_ for seven days prior to nucleofection.

### Next generation sequencing

FastQ files were checked for quality using FastQC (“Babraham Bioinformatics - FastQC A Quality Control tool for High Throughput Sequence Data,” n.d.) and MultiQC (Ewels et al., 2016). Trimming, alignment and quantification of editing was performed with CRISPResso2 v2.2.10 (Clement et al., 2019). The trimmomatic command ILLUMINACLIP:trimmomatic_fastas/TruSeq3-PE-2.fa:2:30:10:2:true MINLEN:75 was used. For base editing samples a quantification center of −15 with a quantification window of 10 was used, whereas a quantification center of −3 and a window of 1 was used. For all samples the minimum homology accepted was 70. For base editor samples the following parameters were added --base_editor_output --conversion_nuc_from A --conversion_nuc_to G. Summary data was compiled from CRISPResso output files and tabulated for downstream analysis. Default parameters were used with the genomic region and gRNA sequence for each sample.

### Subretinal administration in wild type mice

Studies were performed at either Pharmalegacy or Powered Research with the animal study and procedure reviewed and approved by the Pharmalegacy or Powered Research IACUC, respectively. For studies performed at Pharmalegacy 6- to 8-week-old male C57BL/6J were acclimatized for a minimum of 3 days and enrolled into the study. Animals were examined using a slit lamp and indirect ophthalmoscope with animals with no abnormal findings enrolled randomly into the study groups. Animals were anesthetized with Zoletil (25–50 mg/kg, intraperitoneally (IP)) + Xylazine (5 mg/kg, IP). The ocular surface was disinfected with iodophor, the sclera of the inner side of the corneal and scleral limbusa punctured with a 30G disposable injection needle, and 1 μl of test article administered via a microinjector with a 34G flat needle (utilizing the puncture port to enter, bypassing the lens to reach the vitreous, avoiding main blood vessels, with gradual insertion of the needle into the subretinal space with a slow bolus). Immediately after the syringe was removed the needle port was compressed with a cotton swab for 5 seconds. The injection procedure was then repeated on the contralateral eye. Optical coherence tomography (OCT) was then utilized to confirm whether the injection was successful (i.e., formation of a subretinal bleb). For post-injection animal care, eyes were treated with levofloxacin eye drops and ofloxacin eye ointment once in the morning and once in the afternoon for three consecutive days post dosing in addition to continued body weight measurement and clinical observation. 7 days post dosing animals will be euthanized via CO_2_ inhalation followed by cervical dislocation. Eyes collected for gDNA extraction were dissected immediately to separate retina from the remaining eyecup. The retina was placed in DNase/RNase free tubes and snap frozen in liquid nitrogen and stored in - 80 °C. The remaining eye cup was transferred to pre-cooled DMEM supplemented with 10 % FBS and then RPE cells manually scraped off from choroid/sclera and collected via centrifugation. Genomic DNA was then extracted from the resulting cell pellet for subsequent analysis.

For studies performed at Powered Research, 6–8-week-old female C57Bl/6J were acclimated and then enrolled on to study. Animals were given buprenorphine 0.01–0.05 mg/kg subcutaneously (SQ) and a cocktail of 1.0% tropicamide HCl and 2.5% phenylephrine HCl (Tropi-Phen) topically to dilate and proptose the eyes. Animals were tranquilized for the surgical procedure with a ketamine/xylazine cocktail (80–90/10–20 mg/kg) administered IP, and one drop of 0.5% proparacaine HCl was applied to both eyes. The cornea was kept moistened using topical eyewash, and body temperature was maintained using a heated surgical table and hot pads as needed. A small pilot hole was made just superior to the limbus of the eye near the 12 o’clock position using the tip of a 30G needle. The subretinal (SR) injection was made using a 34G needle and a 10 µl syringe (World Precision Instruments). This procedure was repeated on the contralateral eye. Following injections, animals underwent OCT imaging to confirm dose location. If at any time during the surgical procedure or following OCT scanning, the surgeon determined the injection was suboptimal or not successful (as defined by the inability to see a subretinal bleb by OCT imaging), the animal was euthanized and replaced. Following the procedure, 1 drop of neomycin polymyxin B sulfates gramicidin ophthalmic solution or ofloxacin was applied, followed by eye lubricant. Mice were given atipamezole to reverse the xylazine effects (0.1-1.0 mg/kg) and were allowed to recover normally. Cage-side clinical observations were performed once daily, with particular attention paid to the eyes. All observations were recorded. All animals were observed twice daily by a technician for mortality, abnormalities, and signs of pain or distress.

While remaining sedated from Day 8 OCT procedures or while sedated as previously described, all animals were humanely euthanized via carbon dioxide asphyxiation and death was confirmed by cervical dislocation or thoracotomy.

Immediately following euthanasia, all enrolled eyes were enucleated and placed into a 24-well dish containing phosphate-buffered saline (PBS). Under a dissecting scope, extraneous muscles and connective tissue were gently removed from the back of the eye. The anterior segment components (cornea, lens) were removed and discarded. Then, the retina was pulled away from the RPE/choroid/sclera and collected into a pre-weighed 2 ml Precellys homogenization tube and re-weighed. The remaining RPE/choroid/sclera was then collected into a pre-weighed 2 ml Precellys homogenization tube and re-weighed. Samples were stored at −80 °C until further processing.

Immediately following euthanasia, all enrolled eyes were enucleated and placed into a 24-well dish containing phosphate-buffered saline (PBS). Under a dissecting scope, extraneous muscles and connective tissue were gently removed from the back of the eye. The anterior segment components (cornea, lens) were removed and discarded. Then, the retina was pulled away from the RPE/choroid/sclera and collected into a pre-weighed 2 ml Precellys homogenization tube and re-weighed. The remaining RPE/choroid/sclera was then collected into a pre-weighed 2 ml Precellys homogenization tube and re-weighed. Samples were stored at −80°C until further processing.

To isolate genomic DNA, the tissues were thawed on ice and homogenized, and DNA was isolated using the Quick-DNA™ Miniprep kit (Zymo, Cat# D3025) following the manufacturer’s instructions for solid tissue samples. Briefly, RPE/choroid and retina samples were incubated in Lysis Buffer with or without Proteinase K, respectively, and homogenized to solubilize the tissue. Next, the tissue lysate was spun through a Zymo-SpinTM IICR Column. The column was washed with DNA Pre-Wash Buffer, then washed with g-DNA Wash Buffer. Finally, DNA was eluted in DNA elution buffer. DNA integrity and concentration was dually evaluated using a NanoDrop 1000 spectrophotometer and agarose gel electrophoresis. Based on the Zymo kit information, a yield of between 1-3 µg DNA/mg of tissue was expected. All samples passed the integrity QC with a 260/280 value between 1.86 and 2.00 and the agarose gel showed a bright, single band above 1 kb for all samples. The average DNA yield for retina was 3.8 µg/mg and the average yield from the RPE/choroid/sclera was 1.1 µg/mg.

### Subretinal injection in Ai14 reporter mice

All mice were handled in accordance with the guidelines and protocols provided by the Association for Research in Vision and Ophthalmology Statement for the use of animals in ophthalmic and vision research and were approved by Institutional Animal care and use committee of University of Wisconsin-Madison. Ai14 reporter mice of either sex from Jackson Laboratory were used. 4–6-week-old adult mice were anesthetized with IP injection of Ketamine (80 mg/kg) / Xylazine (16 mg/kg) cocktail. Body temperature of anesthetized mice was maintained by keeping mice on a heating pad throughout the surgery process until the mouse was awake. The eyes were dilated with 1 % tropicamide, and a topical anesthetic, 0.5% proparacaine hydrochloride, was applied to the eyes before the surgery. The eRNP complex was injected to the subretinal space using a 34G blunt needle connected to a UMP3 nanofil microsyringe pump that controls the volume and the speed of the injection. 2 µl of the solution containing the eRNP complex was injected and the needle was left at the site of the injection for additional 10-15 secs before withdrawing.

To assess the tdTomato expression generated by successful gene editing, the mice were sacrificed and their eyes were collected 2 weeks after injection. Enucleated eyes were cleaned twice with PBS and a puncture was made using an 18G needle at ora serrata. The corneal incisions were used to open the eyes, and the lens was carefully removed. To flatten the RPE and get the final “starfish” appearance, incisions were made radially to the center of the eye cup. The retina was then gently separated from the RPE layer. The separated RPE was flat-mounted on the cover-glass slide and imaged with NIS-Elements using a Nikon C2 confocal microscope (Nikon Instruments Inc.) and ImageJ software (NIH) was used for analysis.

### Subretinal administration in wild-type minipigs

On the day prior to subretinal dosing, animals received 5 mg/kg methylprednisolone intramuscularly (IM). On the day following surgery and twice daily (BID) for 2 additional days, animals received 1 drop of NeoPolyDex topically OU. Animals were fasted the morning prior to the procedure. On Day 1, animals received a topical mydriatic agent (1.0% tropicamide HCl) at least 15 minutes prior to anesthesia. Buprenorphine (0.03 mg/kg) and atropine (0.05 mg/kg) were administered IM. Animals were anesthetized with a ketamine/dexmedetomidine cocktail (10– 15/0.05–0.3 mg/kg IM).

The area around the eyes including the eyelids was cleaned with a 10% solution of baby shampoo using gauze. The shampoo was rinsed with sterile saline, and a 5% betadine solution was applied topically followed by a rinse with sterile saline. The animal was then transferred to the surgical suite. Using a sterile cotton-tipped applicator soaked in 1% betadine, the conjunctival cul-de-sacs and under the third eyelid were swabbed followed by rinsing with sterile saline. Then the animal received one drop each of 0.5% proparacaine HCl and 10.0% phenylephrine HCl topically in both eyes. A wire eyelid speculum was placed, and a 1.2 mm keratome was used as needed to create an incision through the limbus and sclera-pars plana to reduce intraocular pressure. Wet field cautery was used to prepare the sites intended for the insertion of 23G ports (trocar-cannula entry systems), 3–4 mm posterior to the superior limbus. A local retinal detachment was created in the superior region of the eye, centered on the area centralis at the mid-line. The injection was made using a 38G extendable polytip cannula SR injection needle (MedOne, Sarasota, FL) at a rate of ∼100 µl over a 5 second push. The conjunctiva was closed with suture if needed. Following injection procedures, NeoPolyDex was applied topically. The procedure was repeated on the contralateral eye. Following bilateral injections, animals underwent post-dose OCT imaging to confirm the presence of an SR bleb. Animals then receive atipamezole (0.5 mg/kg IM) to reverse the effects of dexmedetomidine and were allowed to recover normally. Approximately 8–12 hours following surgery, animals received a second dose of buprenorphine (0.3 mg/kg IM).

Cage-side clinical observations occurred once daily, with particular attention paid to the eyes. All observations were recorded. All animals were observed twice daily for mortality, abnormalities, and signs of pain or distress. Body weights were taken at baseline, weekly, and at the time of necropsy. Ocular examinations were performed using a slit lamp biomicroscope and indirect ophthalmoscope to evaluate toxicologic or inflammatory changes of the ocular adnexa, cornea, and anterior and posterior segments of the eye. The semiquantitative preclinical ocular toxicology scoring (SPOTS) system was used for scoring. A topical mydriatic (1% tropicamide HCl) was administered prior to ocular examinations to facilitate examination of the posterior segment. Animals were not tranquilized for examinations.

### Subretinal administration in SRM-1 reporter pigs

SRM-1 transgenic reporter animals (Recombinectics, SRM-1) were administered 5 mg/kg methylprednisolone intramuscularly (IM) one day prior to injections, as well as one drop of neomycin polymyxin B sulfates dexamethasone (NeoPolyDex) the afternoon of dosing and twice daily (BID) for two additional days post-surgery. Animals were fasted the morning prior to the procedure. On the day of dosing animals received a topical mydriatic agent (1.0% tropicamide HCl) at least 15 minutes prior to anesthesia. Buprenorphine (0.03 mg/kg) and atropine (0.05 mg/kg) were administered IM. Animals were anesthetized with a ketamine/dexmedetomidine cocktail (10-15/0.05-0.3 mg/kg IM).

eRNP test articles were loaded into a MicroDoseTM Injection Kit (MedOne, #3315) and connected to an extendible PolyTipSR injection cannula (23G×32 mm cannula with 38G, 5 mm length tip (MedOne, 3248)). The MedOne polymeric tip was extended, and the needle was primed with approximately 240 µl of test article. The needle and syringe pair were allowed to rest on an even surface for one minute to equilibrate pressure throughout the system. Then the polymeric tip was wiped of excess solution utilizing sterile gauze, followed by an additional 30 seconds of dwell time. Next, the polymeric tip was retracted, and the needle and syringe were moved to the surgical field. The needle and syringe pair remained horizontal and in the same plane prior to injection. The MicroDose Injector was attached to a Viscous Fluid Control Pak (Alcon, 8065750957) to allow foot pedal-actuated injections at a controlled rate via the Constellation system. Care was taken to avoid excessively kinking or twisting the extension tubing. Prior to injection, if solution was observed at the needle tip, it was carefully wiped with sterile gauze.

Before injection, the area around the eyes including the eyelids was cleaned with a 10% solution of baby shampoo using gauze. The shampoo was then rinsed with sterile saline, and a 5% betadine solution was applied topically followed by a rinse with sterile saline. The animal was then transferred to the surgical suite. Using a sterile cotton-tipped applicator soaked in 1 % betadine, the conjunctival cul-de-sacs and under the third eyelid were swabbed followed by rinsing with sterile saline. Following surgical preparation, the animal received one drop each of 0.5 % proparacaine HCl and 10.0 % phenylephrine HCl topically in both eyes. A wire eyelid speculum was placed, and a 1.2 mm keratome was used as needed to create an incision through the limbus and sclera-pars plana to reduce intraocular pressure. Wet field cautery was used to prepare the sites intended for the insertion of 23G ports (trocar-cannula entry systems), 3–4 mm posterior to the superior limbus. A local retinal detachment was created in the superior region of the eye, centered on the area centralis at the mid-line. The test article was administered using the primed MedOne injector via foot pedal-actuation with the Constellation System using a pressure of 9–20 PSI (lower end for saline, higher end for a gel) at a rate of approximately 100 µl over 5 seconds. The conjunctiva was closed with suture if needed. Following injection procedures, NeoPolyDex ophthalmic solution was applied topically. The procedure was repeated on the contralateral eye. Following injection procedures of both eyes, animals will undergo post-dose OCT imaging to confirm and SR bleb. Animals received atipamezole (0.5 mg/kg) IM to reverse the effects of dexmedetomidine and were to recover normally. In the afternoon of dosing and BID for 2 additional days, animals received 1 topical drop of NeoPolyDex OU, and 8-12 hours after surgery, a second dose of buprenorphine (0.03 mg/kg IM) was administered.

Cage-side clinical observations occurred once daily, with particular attention paid to the eyes. All observations were recorded. All animals were observed twice daily for mortality, abnormalities, and signs of pain or distress. Body weights were taken at baseline, weekly, and at the time of necropsy. Ocular examinations were performed using a slit lamp biomicroscope and indirect ophthalmoscope to evaluate toxicologic or inflammatory changes of the ocular adnexa, cornea, and anterior and posterior segments of the eye. The semiquantitative preclinical ocular toxicology scoring (SPOTS) system was used for scoring. A topical mydriatic (1% tropicamide HCl) was administered prior to OEs to facilitate examination of the posterior segment. Animals were not tranquilized for examinations.

### Optical coherence tomography (OCT)

Animals were fasted the morning of anesthetic administration unless the procedure was terminal. animals were administered atropine (0.05 mg/kg IM), then anesthetized with ketamine/dexmedetomidine (10–15/0.05–0.3 mg/kg IM). One drop of 1% tropicamide HCl and 0.5% proparacaine HCl was applied to both eyes. OCT was performed using a Spectralis HRA OCT II (Heidelberg Engineering). A wire eyelid speculum was placed as needed and/or forceps were used to raise the periocular tissue, and eyewash may have been applied topically. A series of b-scans was acquired using a 55° lens to capture the treated area (within the bleb) and the untreated area in the inferior region. Following completion of OCT procedures, animals underwent fundus imaging. Three measurements of the outer nuclear layer (ONL) thickness and total retinal thickness (TRT) were made from a b-scan in the treated area (within the bleb) and untreated area (in the inferior region).

### Histological analysis of porcine eyes

Immediately following euthanasia, eyes were enucleated and the 12 o’clock position of the eyes was marked; the whole eye with attached optic nerve was preserved in Davidson’s fixative for 24 hours and then transferred to 70% ethanol. Eyes were slightly trimmed sagittally on both sides, edge trimmings were discarded. Eyes were then trimmed sagittally in the plane from 12 o’clock to 6 o’clock through the lens and optic nerve along the midline; a photograph was taken of each eye after bisection through the midline to document the orientation/position of the cut. Each half of the trimmed globe was embedded in its own cassette (Cassette A is the nasal half of the eye, Cassette B is the temporal half of the eye). The sectioning approach was informed by the post dose OCT and color fundus images. Ten (10) slides/level at 4 levels (750 µm between levels) were collected from each block, nasal and temporal, for a total of 8 levels (80 slides) total per eye. The first level was as close to midline as possible to capture the bleb region. Eight (8) slides per eye (1 slide/level) were stained with H&E.

### Cas9 immunohistochemistry

Cas9 IHC was performed at Ensigna Biosystems. FFPE sections were sectioned (tissue collected 1 hour post dosing) and dewaxed. Antigen retrieval was performed using pH 6.2 HIER in a Biocare Medical MACH4 HRP polymer block for 5 minutes at room temperature (Biocare Medical MRH534L). Primary detection antibodies utilized were anti-Cas9 rabbit mAb (Clone E7MIH, Cell Signaling Technologies) at 1:100 (0.59 µg/ml) and anti-rabbit IgG (Cell Signaling Technologies) at 1:100 (25 µg/ml). Primary antibodies were incubated for 45 minutes at room temperature, washed with TBS Automation Wash Buffer at 1× (Biocare Medical TWB945M), and then developed at room temperature for 20 minutes with Vina Green (Biocare Medical BRR807AS). Slides were then washed for 10 minutes at room temperature in DI water, stained with hematoxylin for 30 seconds, bluing solution in room temperature DI water, dehydrated, and then coverslipped. The ONCORE Pro automated platform from Biocare Medical was used to run the IHC on the porcine eye samples.

### Immunofluorescence imaging of cryosections

Eyes selected for IF were enucleated and marked with metallic Sharpie superiorly at the 12 o’clock position. A 3 mm incision was made at the limbus and then the eye was placed in 4% paraformaldehyde (PFA) in phosphate-buffered saline (PBS) for 24 hours at RT. The anterior segment and lens were dissected away, and vitreous humor (VH) was gently removed. Eyecups were brought through a sequential sucrose gradient (10–30%, 1 hour each) followed by embedding in OCT medium oriented such that the superior surface was toward the embedder. Once tissue was positioned, blocks were frozen on dry ice and held at −80°C until sectioning.

To survey the dosed region of the pig eye, blocks were sectioned (14 μm) at 3 levels across the eye. The surface closest to the bleb (nasal or temporal) was sectioned first, and sets of contiguous ribbons of sections (n = 3–4 sections/slide) were captured onto two slides every 200–300 µm at the following 3 levels, using Day 1 post-dose OCT and bleb relation to the optic nerve head (ONH) to gRNA: just temporal to the bleb (10 slides), within the bleb (∼25–30 slides), and just nasal to the bleb (10 slides).

Slides were blocked (PBS with 0.05 % Triton X-100, 2 % Bovine Serum Albumin [BSA] and 2 % normal donkey serum [NDS]) and then incubated overnight at 4 °C in 1:100 goat anti-tdTomato (LS Bio, LS-C340696) and 1:100 mouse anti-RPE65 (Novus Biologicals, NB100-355). Following three 5 minute PBS washes, slides were then incubated for 2 hours at room temperature with corresponding conjugated secondaries (1:200 donkey anti-goat Cy5 [Jackson ImmunoResearch 705-175-147], and 1:200 donkey anti-mouse Cy2 [Jackson ImmunoResearch 715-545-150]) and 1:1000 DAPI. tdTomato was detected via direct fluorescence (native signal) and indirect detection. As a negative, control an additional slide was stained with anti-goat IgG isotype control (ThermoFisher Scientific 31245), 1:100 mouse anti-RPE65 (Novus Biologicals NB100-355), and 1:1000 DAPI.

Following 3 final washes, slides were mounted and coverslipped. Slides were imaged on an Olympus Bx63 upright fluorescent microscope equipped with plan apochromatic objectives and Olympus CellSens software. For each eye, a 4× tiled overview and a 20× higher magnification image were captured where for the region where tdTomato signal was the broadest and brightest.

### Flat-mount imaging

Eyes utilized for flat-mount analysis were enucleated with a tail of ON attached, and a #11 blade was used to make an incision at the border of the clear cornea and sclera, taking care not to cut through the ciliary body. Curved scissors were used to cut away the anterior segment, leaving the retina attached at the level of the ciliary body. VH was poured out of the remaining eyecup and eyecups were fixed for 24 hours in freshly prepared 4 % PFA at 4 °C. The following day, eyecups were transferred to 1× PBS, the back of the eye was thoroughly cleaned of any extraneous tissue, the ciliary body was trimmed away, and the retina was carefully isolated. The RPE/choroid and retina were then washed 3×5 minutes in PBS containing 0.1 % Triton X-100 (wash buffer) and both the RPE/choroid and retina were blocked for 2 hours at room temperature blocking with PBS + 5 % NDS and then stained overnight at room temperature with 1:100 anti-tdTomato (LS Bio, LS-C340696).

Following incubation in secondary antibody (1:200 donkey anti-goat Cy5, Jackson ImmunoResearch 705-175-147), RPE/choroids and retinas were washed 5×5 minutes in wash buffer. An imaging spacer was used to create a chamber on the slide. Radial cuts were then made toward the center to allow the tissue to lay flat, and the tissues were mounted in FluoroGel and coverslipped. The retina was mounted such that the photoreceptor layer was against the coverslip. A 10× tiled overview of the region of expression and a 20× high magnification in the region of the brightest, broadest tdTomato signal were captured using an Olympus Bx63 upright fluorescent microscope equipped with plan apochromatic objectives and CellSens software. Exposure settings were set based on the brightest signal in the targeting gRNA sample to ensure the signal was not oversaturated. Exposure settings were held constant across Groups.

---

**Supp Figure 1.**
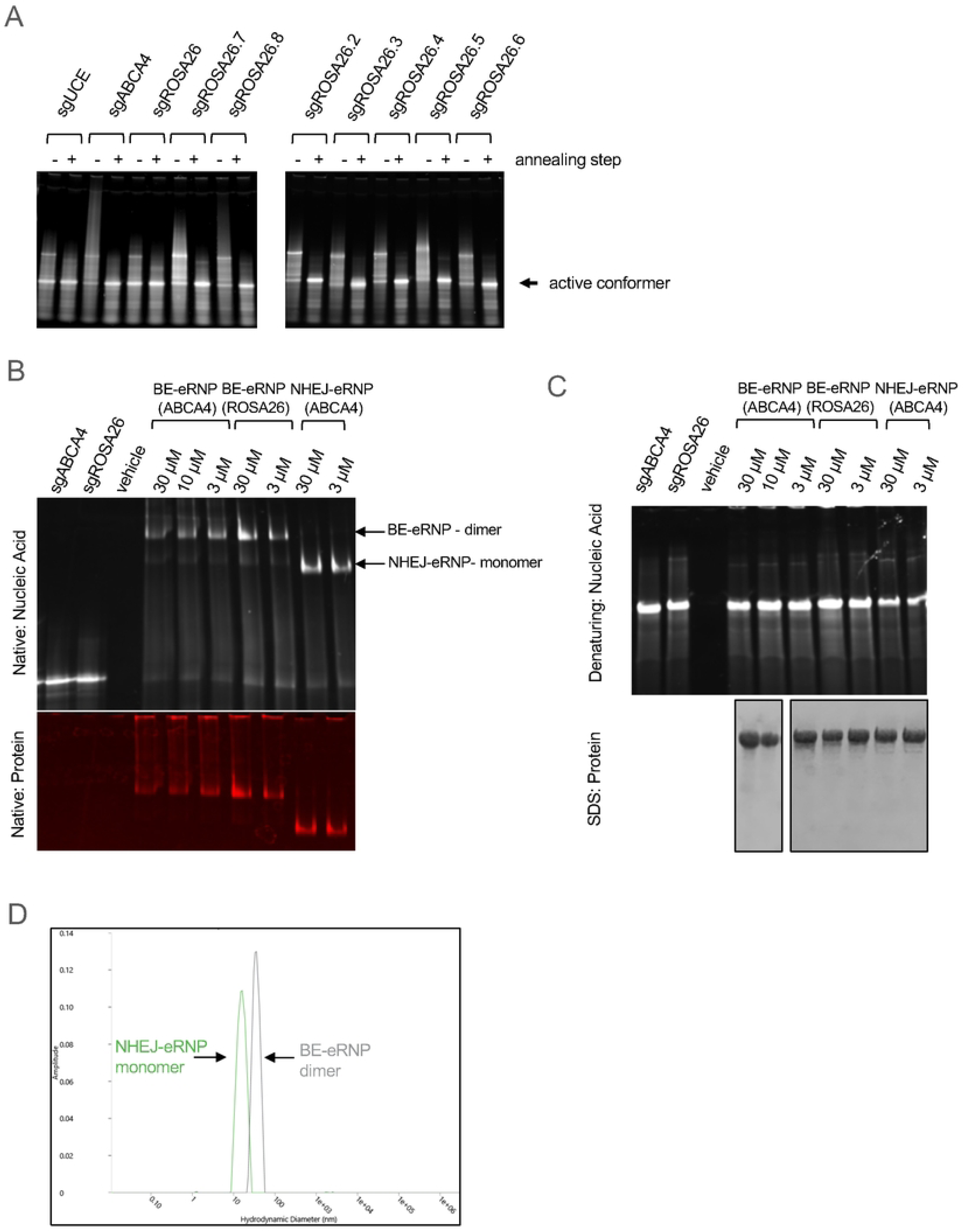
Analysis of gRNA, Cas nucleases, and RNPs prior to subretinal injection in minipigs. **A.** Native gel electrophoresis of sgRNAs prior to eRNP assembly to evaluate RNA conformation**. B.** Native gel electrophoresis of assembled NHEJ-eRNPs and BE-eRNPs to determine quaternary structure. **C.** Denaturing PAGE and SDS-PAGE of NHEJ-eRNPs and BE-eRNPs to analyze purity of total RNA and protein. **D.** Dynamic light scattering of NHEJ-eRNP and BE-eRNP. gRNA annotations are described in Supp Table 2.

**Supp Figure 2.**
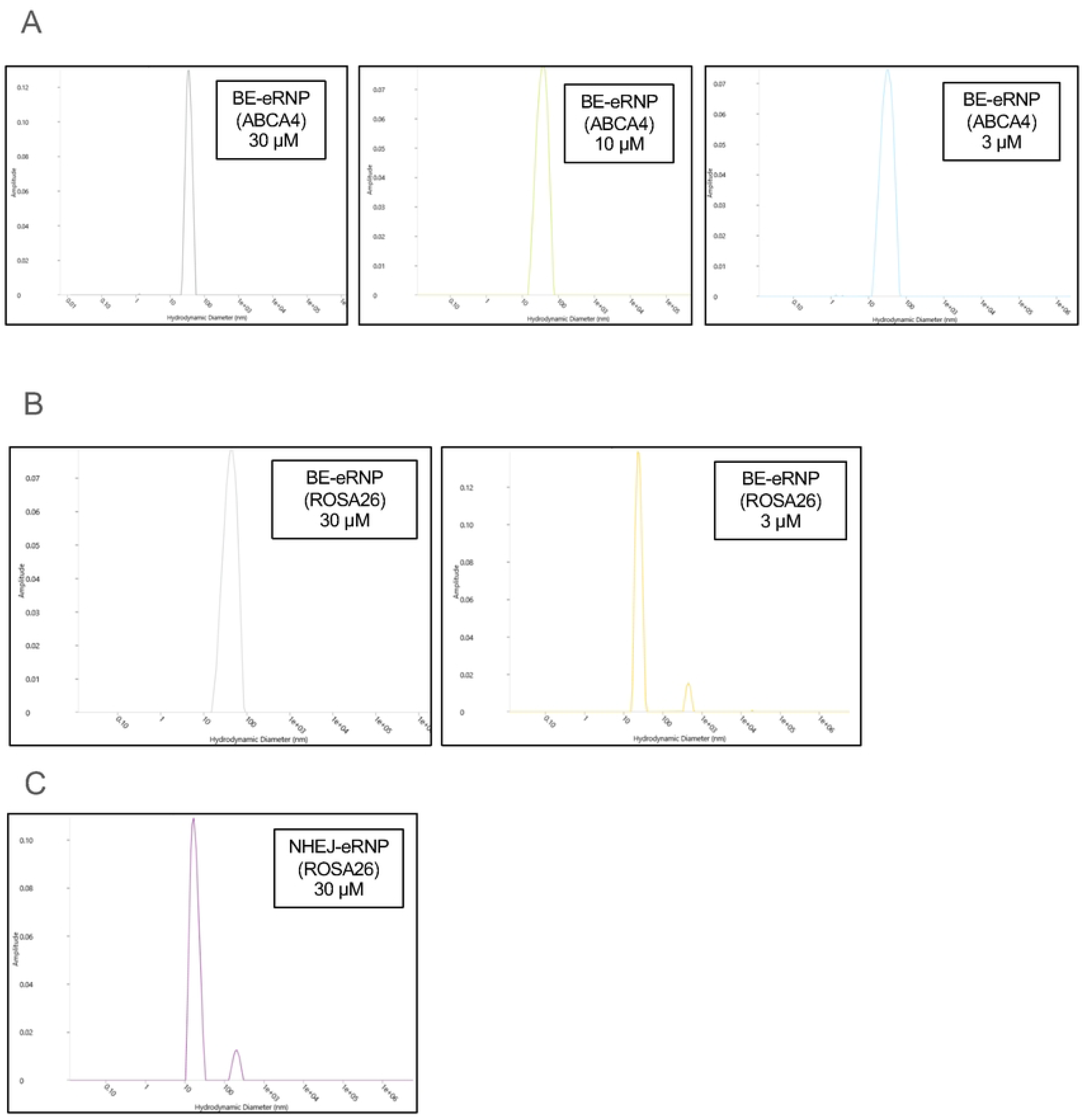
Dynamic light scattering data for all test articles. **A.** BE-eRNP (assembled with sgRNA targeting Abca4 locus) at 3 different dosing concentrations. **B.** BE-eRNP (assembled with sgRNA targeting the Rosa26 locus) at 2 different dosing concentrations. **C.** NHEJ-eRNP (assembled with sgRNA targeting the Rosa26 locus) at a single dosing concentration.

**Supp Figure 3.**
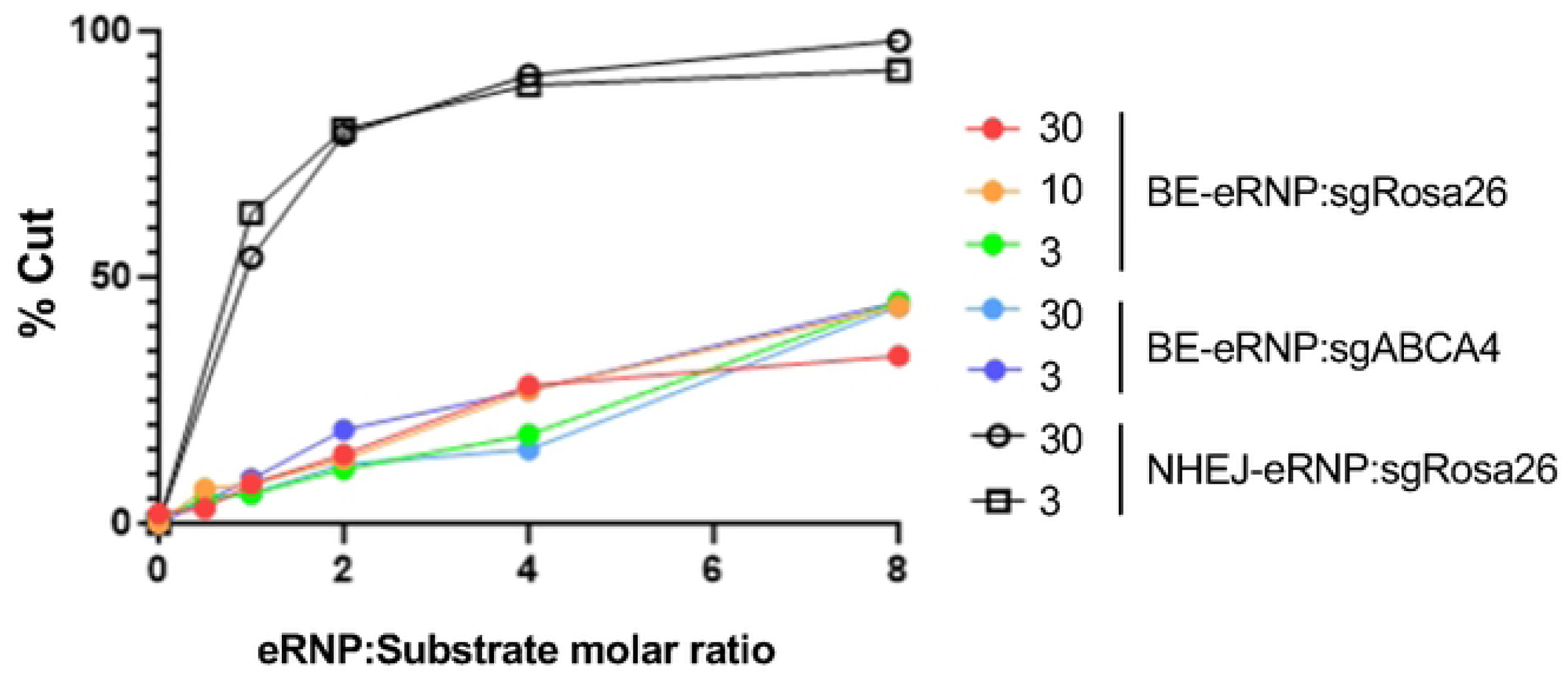
Enzymatic activity of NHEJ-eRNPs and BE-eRNPs. In vitro DNA cleavage (NHEJ-eRNPs) and adenosine deamination (BE-eRNPs) assay data are displayed.

**Supp Figure 4.**
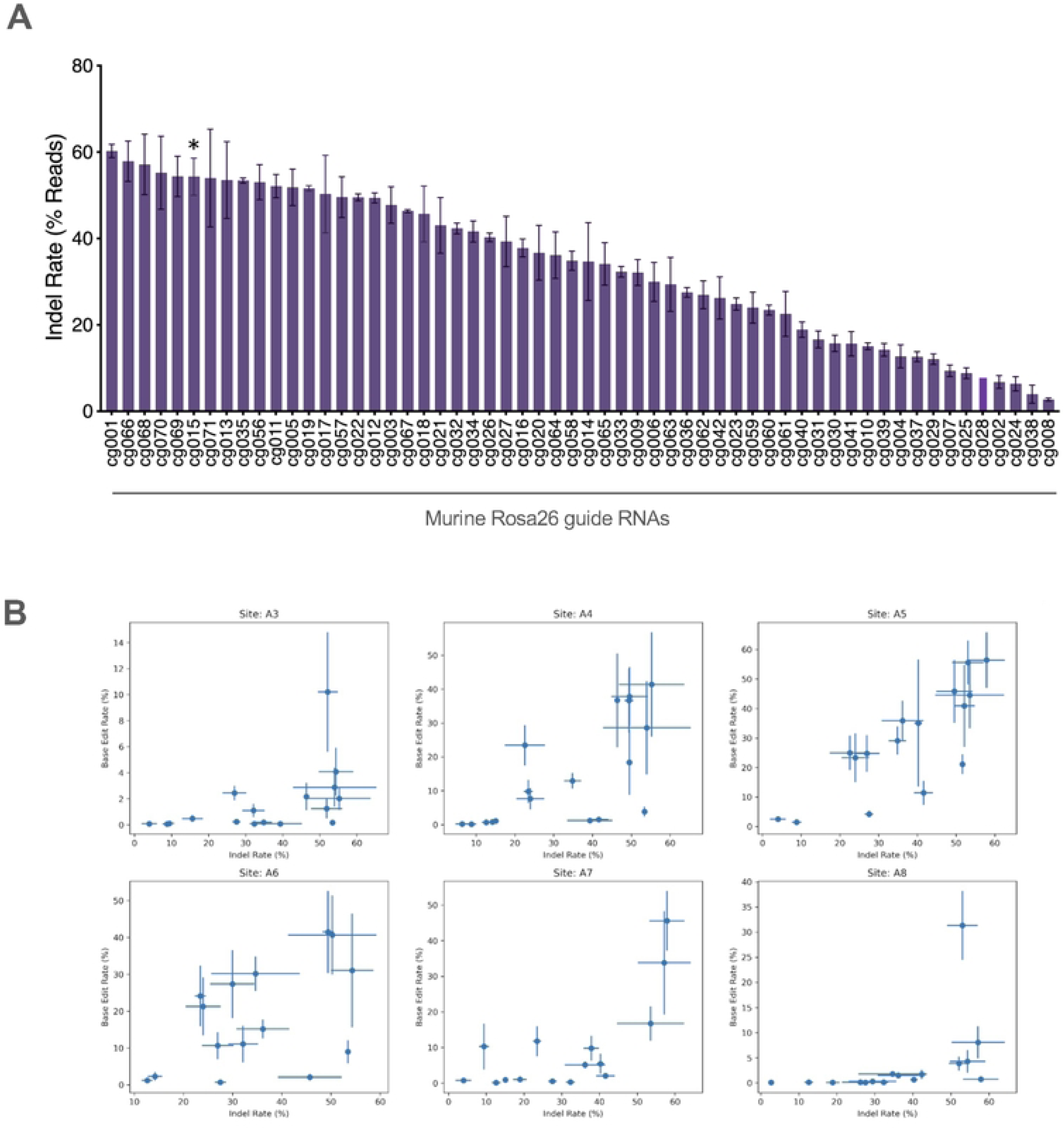
Identification of high efficiency NHEJ gRNAs targeting the mouse *Rosa26* locus and site-wise comparison of adenine base editing rates versus indel rates. **A.** Mouse T cell editing rates versus gRNA for NHEJ-eRNPs. NHEJ-eRNPs were complexed with various gRNAs targeting the *Rosa26* locus and then used to treat stimulated mouse T cells. NHEJ-eRNPs were complexed with the indicated gRNAs in crRNA:trcrRNA format and then nucleofected into cells. Editing rates were quantified based on the frequency of reads containing indels from Illumina sequencing. The asterisk indicates the crRNA corresponding to the targeting gRNA used in main text Fig. 2B and 2C. **B.** Frequency of A→G transitions upon ABE-eRNP treatment vs. frequency of indels upon NHEJ-ABE treatment at each A position across protospacers tested in Fig. 2A and panel A of this figure. For the A position indicated in each panel, the ABE and indel rates for subset of gRNAs with an A at that position were plotted against each other. Each point corresponds to the mean editing rate for one gRNA and error bars represent the standard deviation across biological replicates.

**Supp Figure 5.**
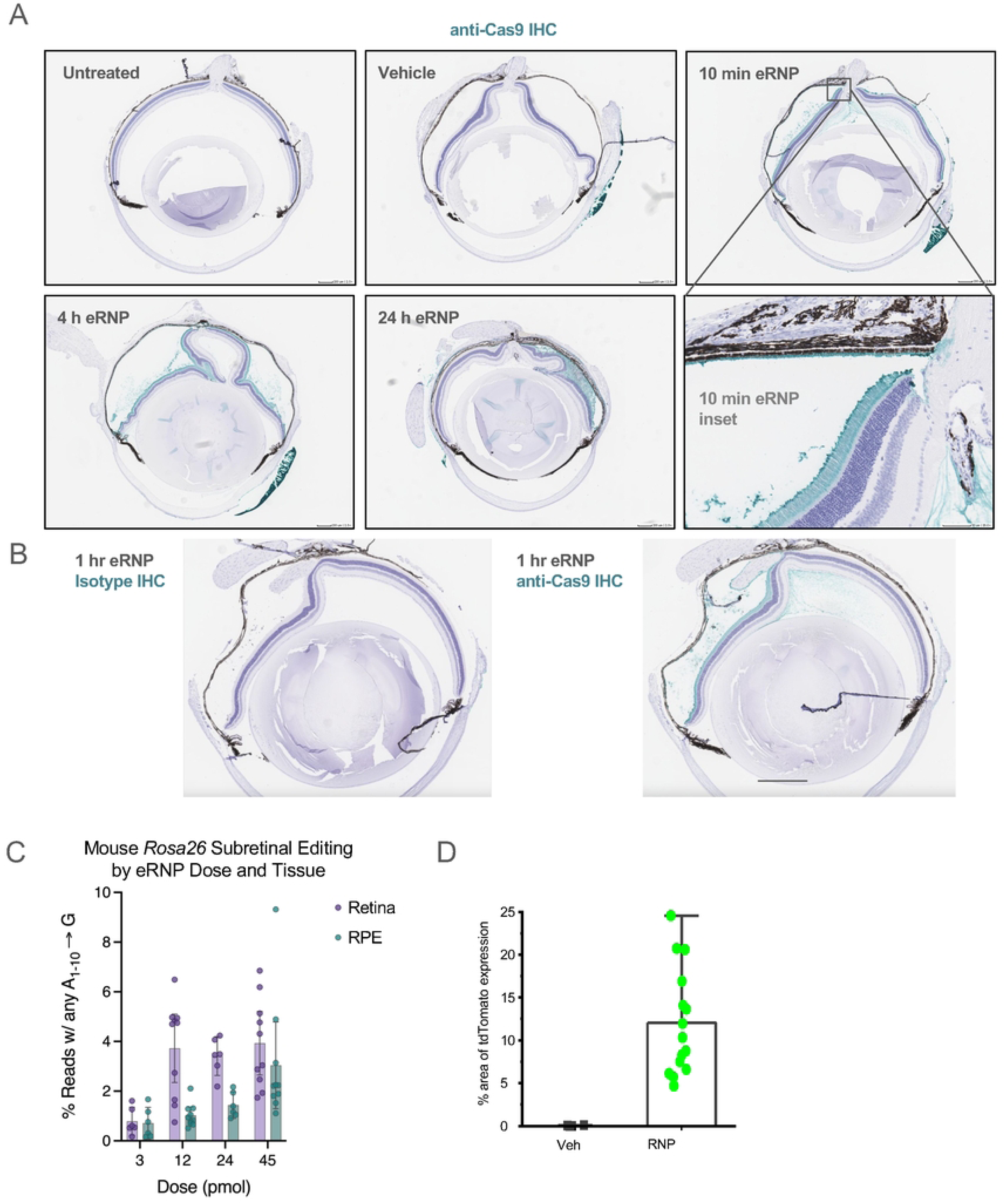
**A.** Representative images of murine eyes treated with subretinal administration of 2 µl of vehicle or eRNP at denoted time points with an untreated eye control. Sections were stained by anti-Cas9 immunohistochemistry (teal). **B.** Representative images of murine eyes 1 hour post subretinal administration with 1 µl eRNP stained with isotype control (left) or anti-Cas9 immunohistochemistry (right, teal) **C.** Editing rates by tissue following subretinal injection of *Rosa26*-targeting ABE-eRNPs complexed with mmRosa_sg1 in mice at four doses. Five days following subretinal administration, mice were sacrificed and their eyes dissected to separate the neural retina from the eyecup, and then RPE layer was separated from the choroid. Editing rates from Illumina DNA amplicon sequencing are shown as mean ± SD across eyes at each dose for retina and RPE tissue samples. Each overlaid point corresponds to a single eye. Reads were scored as positive for editing if at least one A→G transition was detected within a 10 base edit window. These data were collected with the same sample shown in Fig. 3 at a different study site and with a different operator.

**Supp Figure 6.**
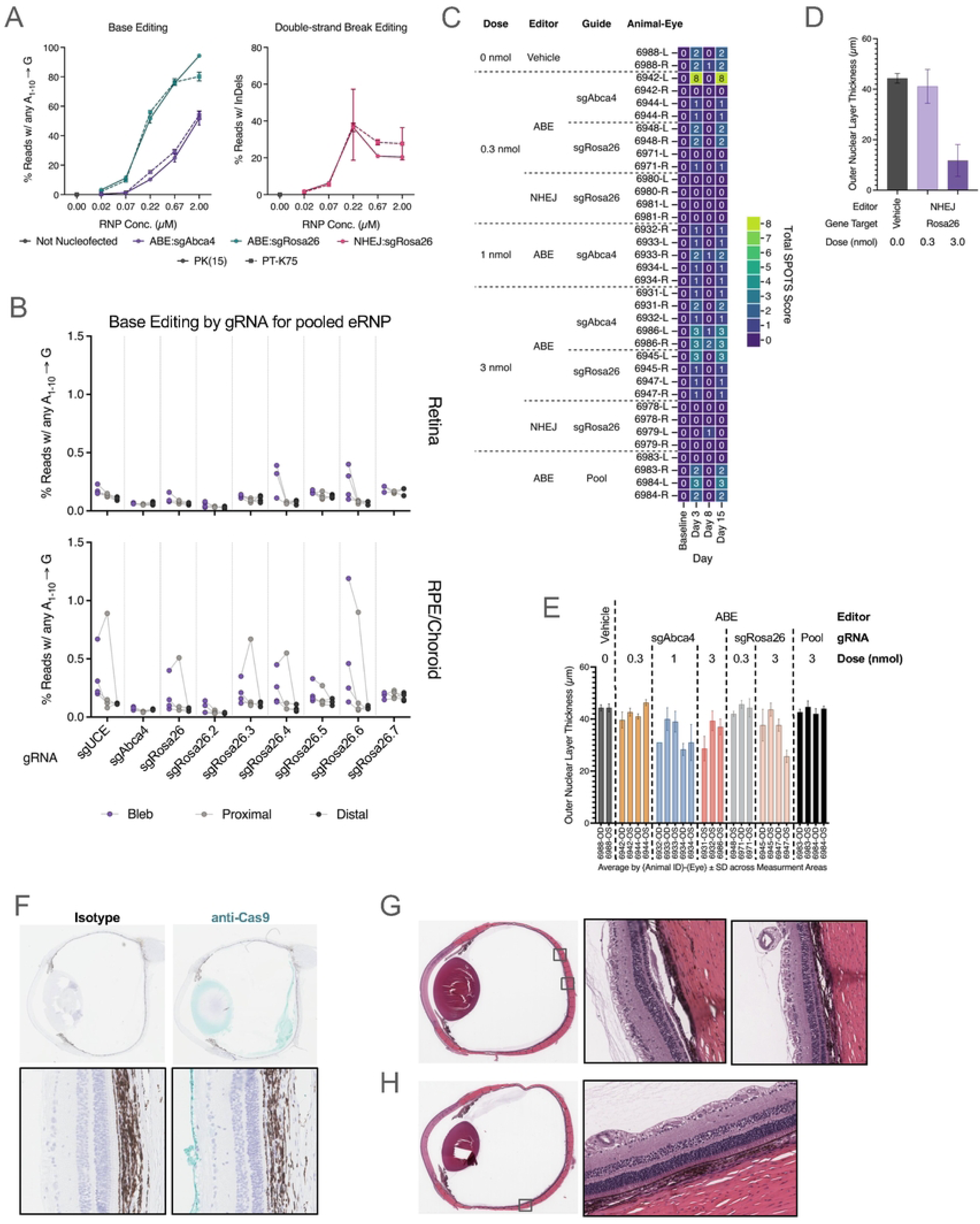
In vitro editing with *S. scrofa*-specific gRNAs and safety-related parameters following eRNP subretinal administration in minipigs. **A.** Adenine base editing and indel formation following nucleofection of BE-eRNPs (left panel) and NHEJ-eRNPs (right panel) in porcine cell lines PK(15) and PT-K75 at various concentrations with two different gRNAs. Editing rates calculated from Illumina DNA amplicon sequencing are plotted as mean ± standard deviation from three biological replicates. The indicated RNP concentrations correspond to the final mixture with cells in the nucleofection cuvette. **B.** Editing rates as shown in (A) and (B) for eyes treated with a pool of ABE-eRNPs each complexed with a different gRNA. Editing rates corresponding to the intended gene target sequence are shown for each gRNA and arranged by tissue. **C.** Aggregate SPOTS scores per minipig porcine eye following subretinal administration of eRNP. **D.** Outer nuclear layer thickness measured by OCT plotted for NHEJ-eRNP groups. Data are plotted as the mean ± standard deviation for each group across all measurements in all eyes. **E.** Outer nuclear layer thickness measured by OCT plotted by eye and grouped by eRNP, gRNA, and dose. Data are plotted as the mean ± SD across measurement areas (regularly spaced 2D OCT slices spanning the superior and inferior regions proximal to the superior injection bleb) collected for each eye. **F.** Representative images of porcine eyes 2 weeks post subretinal administration stained with isotype control or anti-Cas9 immunohistochemistry (teal). Representative whole eye scans and high magnification (inset) images of H&E-stained porcine eyes 2 weeks post subretinal administration with **G**) 3 nmol or **H**) 3 nmol NHEJ eRNP.

**Supp Figure 7.**
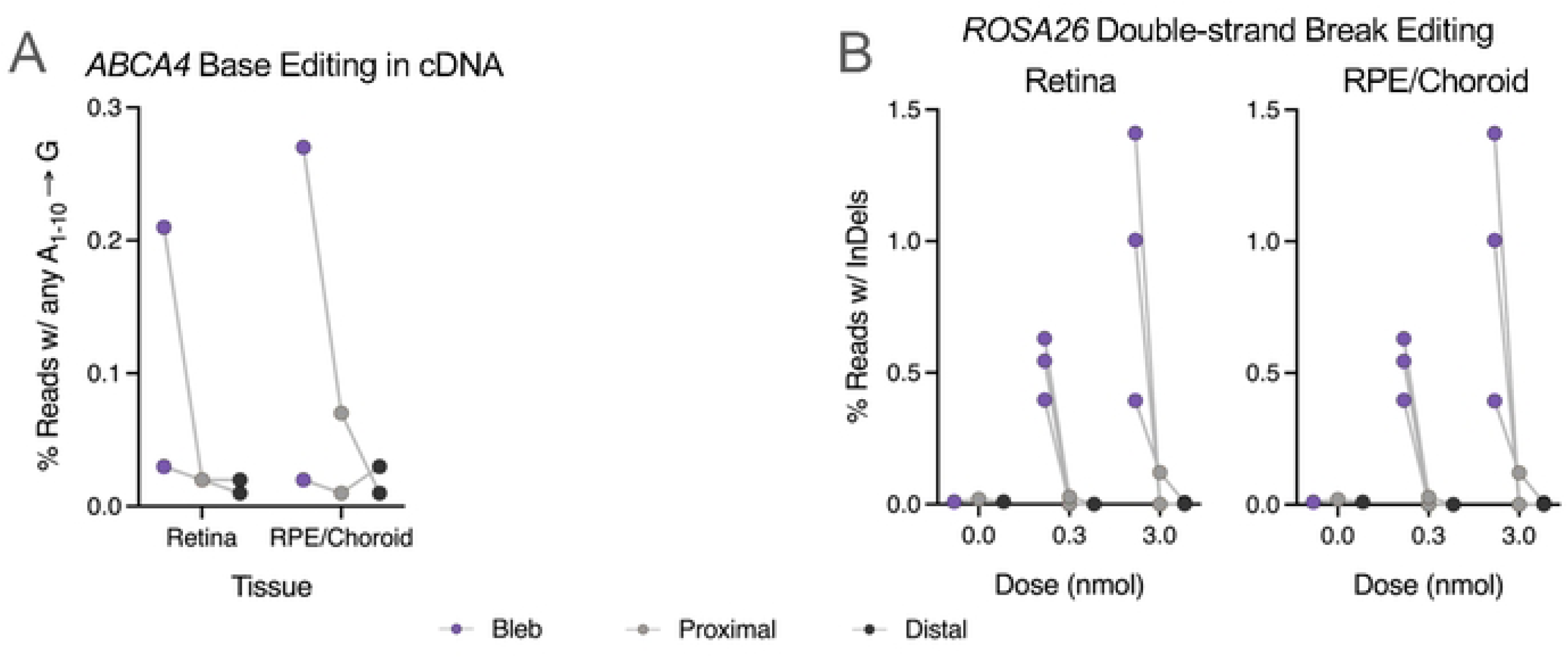
Adenine base editing in cDNA and indel formation following NHEJ-eRNP or BE-eRNP subretinal administration in minipig eyes. **A.** Editing rates by tissue following subretinal injection of *ABCA4-*targeting ABE-eRNPs complexed with sgABCA4 in minipigs. Two weeks following subretinal administration, animals were sacrificed and eyes were dissected to collect intact tissue layers for the neural retina and the choroid + retinal pigmented epithelium. For each tissue, three biopsy punches were taken corresponding to the injection bleb, tissue immediately adjacent to the bleb, and a region distal to the bleb. Editing rates observed in reverse-transcribed cDNA from each sample from Illumina amplicon sequencing are plotted for each eye by tissue and dose as described in Fig. 4. **B.** Editing rates by tissue following subretinal injection of *ROSA26*-targeting NHEJ-eRNPs complexed with sgRosa26 in minipigs at multiple doses. Editing rates from Illumina DNA amplicon sequencing are plotted for each eye by tissue, biopsy region, and dose. Reads were scored as positive if they had an InDel at the expected cut site.

**Supp Figure 8.**
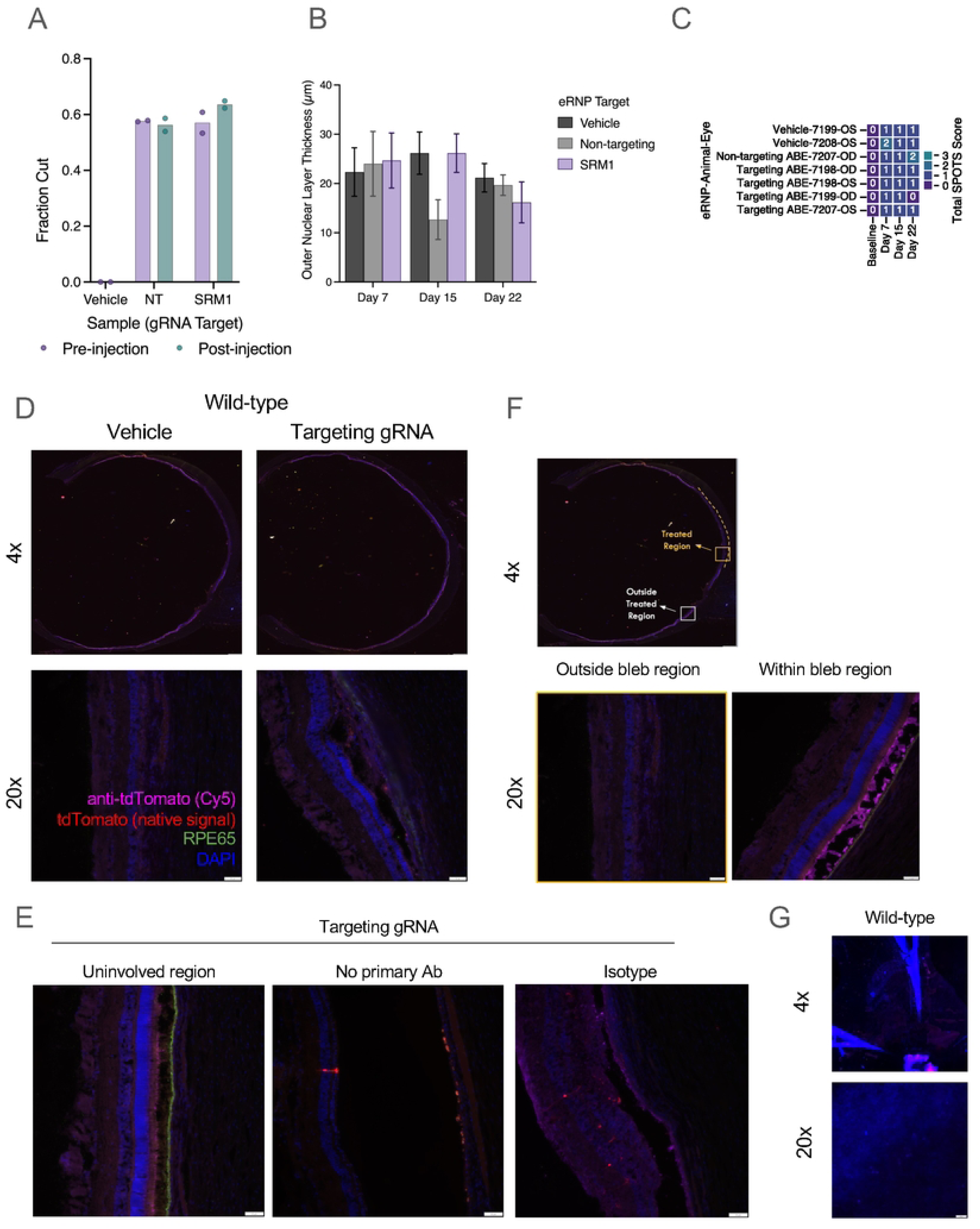
SRM1 eRNP test article characterization and evaluation of activity in SRM1 pigs following subretinal eRNP administration. **A.** Plasmid cutting by purified NHEJ-eRNP samples complexed with non-targeting (NT) and reporter locus gRNAs (SRM1). Sample aliquots were collected and stored before shipment to the test site (pre-injection) and from the material recovered from the injector following administration (post-injection), and then assayed side-by-side. Each point corresponds to a separate reaction. **B.** Outer nuclear layer thickness according to OCT at different time points post injection following NHEJ-eRNP administration in the SRM1 pig study described in Fig. 5. Data are plotted as the mean ± standard deviation for each group across all measurement areas from superior (injected) regions. **C.** Aggregate SPOTS scores per porcine eye following subretinal administration of eRNP. **D.** Fluorescent images of a wild-type littermate (from SRM1 strain) porcine eyes treated with either vehicle or targeting gRNA NHEJ-eRNP. **E.** Fluorescent images from a SRM1 porcine eye from either the uninvolved inferior region, stained without anti-tdTomato primary antibody, or with isotype antibody (corresponding to the anti-tdTomato antibody). **F.** Fluorescent image at 4× and 20× magnification within the treated (superior) and uninvolved (inferior) region of a SRM1 porcine eye. **G.** Fluorescent images of a flat-mounted RPE/Choroid/Sclera from a wild-type littermate control (from SRM1 colony) eye at 4× and 20× magnification.

## Author contributions

Conceptualization - SCW, AJC, MJJ. Methodology, Validation, Investigation, Verification, Formal analysis and Visualization - AJC, JW, RM, KM, AB, PK, SG, JW, CS, RK, GL, SM, VJ, WL, PKS, JC, PB, PO, BGG. Software and Data Curation - KM, PK. Writing (Original draft, review and editing) - SCW, AJC, JW, APS. Writing (Review and editing) - SCW, AJC, BRP, DG, KS, MHF, APS. Supervision - BGG, WL, SCW, AJC, DG, KS, BRP, MHF, MJJ. Project administration and Funding acquisition - MJJ, DG, KS, MHF.

